# Reprogramming insulin receptor activation with a de novo agonist to overcome severe insulin resistance

**DOI:** 10.64898/2026.05.04.722722

**Authors:** Albert Hung, Xinru Wang, Michael Gao, Michelle Ng, Denys Oliinyk, Riley Weaver, Elisabeth Zollbrecht, Sarah Cardoso, Gregory Sebastien Ronel, Chloe Paolucci, Jianxiang Ye, Qing R. Fan, Elizabeth Rhea, Matthias Mann, David Baker, Domenico Accili, Eunhee Choi

## Abstract

Computational protein engineering provides a powerful approach to address longstanding clinical challenges. Severe insulin resistance syndromes caused by mutations in the insulin receptor (IR) are life-threatening disorders for which effective long-term therapies remain lacking. Here, we define the in vivo activity and therapeutic potential of RF-409, a de novo–designed IR agonist that activates the receptor through a mechanism distinct from insulin. RF-409 exhibits markedly prolonged circulation compared to insulin and produces sustained improvements in glucose homeostasis without detectable adverse effects on body composition or liver function. In a patient-derived IR D707A mouse model of severe insulin resistance, RF-409—but not insulin—activates the mutant receptor, restoring glucose regulation and ameliorating hyperglycemia, hyperinsulinemia, lipoatrophy, and pancreatic atrophy. Mechanistically, RF-409 engages the IR through a noncanonical binding geometry while stabilizing an active conformation resembling that induced by insulin. Phosphoproteomic profiling shows that RF-409 elicits broadly insulin-like signaling with distinct temporal features in receptor-proximal regulation. Together, these findings establish a framework for reactivating dysfunctional receptors and suggest broader applications beyond rare receptoropathies, including diabetes and liver disease.

## INTRODUCTION

The insulin receptor (IR) is a receptor tyrosine kinase that coordinates systemic metabolic homeostasis by coupling insulin binding at the cell surface to intracellular signaling governing glucose uptake, macronutrient storage, and growth control(*1–5*). Precise regulation of IR activation, signaling balance, and trafficking is essential for maintaining metabolic homeostasis across tissues.

Pathogenic mutations in IR disrupt these processes and give rise to severe insulin resistance syndromes, including Donohue syndrome, Rabson–Mendenhall syndrome, and Type A insulin resistance(*6–12*). Many disease-causing variants are missense mutations that impair insulin binding or destabilize receptor activation, resulting in profound defects in downstream signaling despite extreme hyperinsulinemia(*13–16*). Clinically, these disorders manifest as severe metabolic dysregulation, including fasting hypoglycemia, postprandial hyperglycemia, growth retardation, and lipoatrophy, and are associated with high mortality early in life(*17, 18*).

Current therapeutic approaches—including supraphysiologic insulin(*19*), insulin sensitizers(*20*), leptin(*21, 22*), and insulin-like growth factor-1 (IGF-1)(*23–29*)—provide limited metabolic benefit and do not correct the underlying receptor signaling defect. Efforts to directly activate mutant IR using monoclonal antibodies(*30–36*) (e.g., 83-7 and 83-14) have demonstrated partial signaling restoration; however, ligand-induced receptor downregulation markedly limits sustained efficacy. Thus, direct IR activation strategies must contend not only with impaired ligand binding but also with altered receptor fate and signaling durability.

Synthetic IR ligands have offered alternative approaches to bypass defective insulin binding. For example, S597 partially activates insulin-binding–defective IR mutants but elicits a biased signaling output that favors metabolic pathways while failing to reconstitute balanced insulin-like signaling across growth and proliferative axes(*37–40*). Although pathway bias may be advantageous in certain settings, such as in patients with concomitant cancer and diabetes, severe congenital insulin resistance syndromes typically present early in life and are frequently accompanied by impaired growth and developmental abnormalities. In this context, selective activation of metabolic pathways alone may be insufficient, as effective therapy may require more balanced reconstitution of insulin-like signaling across both metabolic and mitogenic programs. More broadly, prior studies have largely focused on acute signaling readouts in cultured cells, often using overexpression systems to assess downstream pathway activation, leaving unresolved how pathogenic IR mutations alter receptor conformational states, signaling architecture, and trafficking behavior under physiological conditions.

## RESULTS

### RF-409 activates IR through a distinct mechanism while mimicking insulin action

To address this challenge, we developed de novo–designed IR agonists using structure-guided computational protein engineering(*41*). RF-409 is an optimized IR agonist selected for enhanced receptor activation and improved pharmacological properties. RF-409 shares no sequence or structural homology with insulin (Fig. 1A). Whereas insulin is a two-chain peptide stabilized by multiple disulfide bonds, RF-409 is a single-chain polypeptide composed of two functional components linked by a rigid α-helix.

**Fig. 1.**
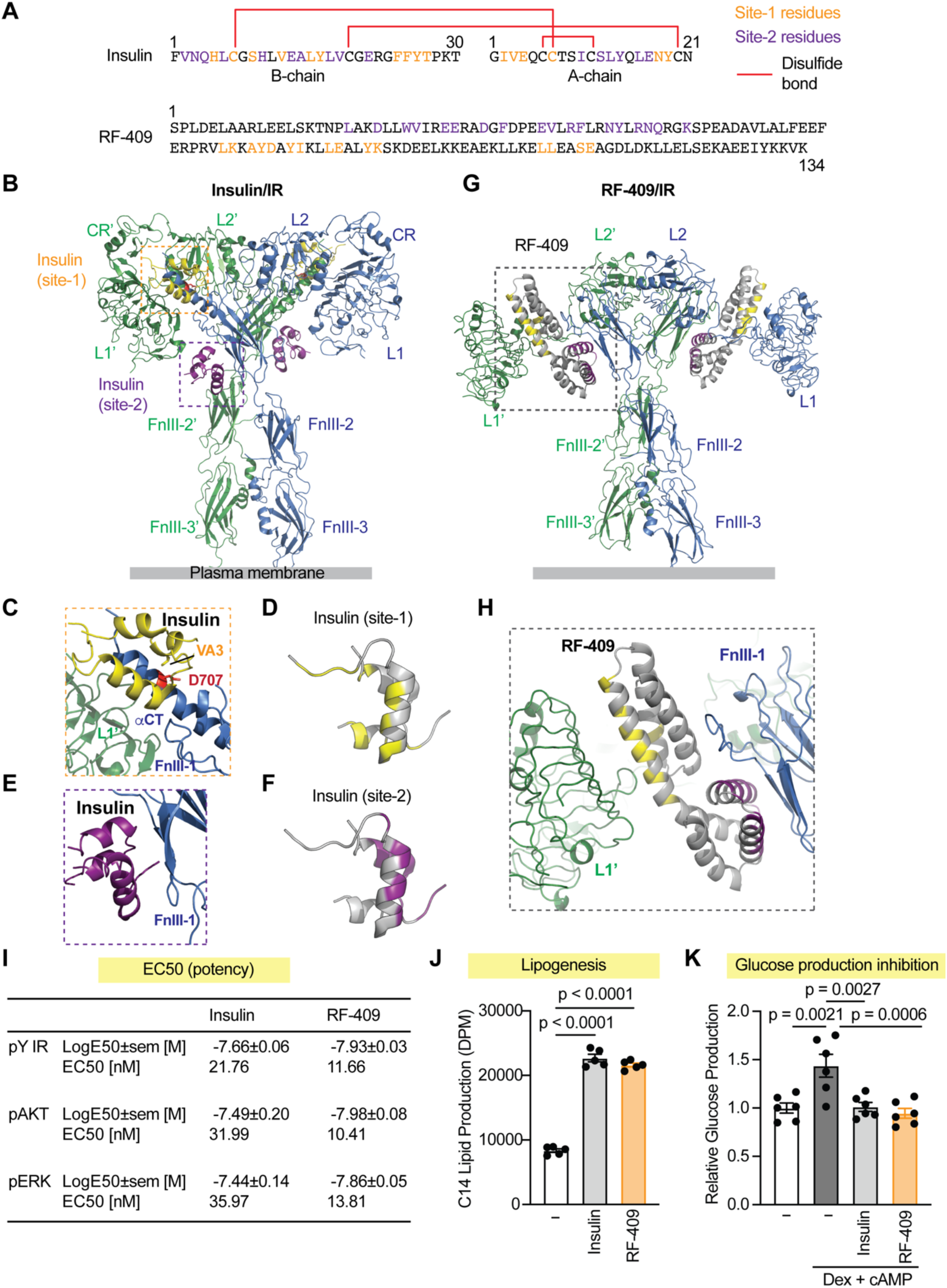
RF-409 is a de novo IR agonist with a distinct binding mode that mimics insulin action. **(A)** Amino acid sequence comparison between insulin and RF-409. Site-1–binding residues are shown in yellow and site-2–binding residues in purple. Disulfide bonds are shown in red. **(B)** Cryo-EM structure of the human IR fully occupied by insulin (PDB: 6PXV). The two protomers are shown in green and blue. Insulin molecules bound at site-1 are shown in yellow, and those bound at site-2 are shown in purple. Asp707 residues on αCT motifs are highlighted in red. **(C)** Structural view of insulin (yellow) bound at site-1 of the IR, engaging the L1 domain (green) from one protomer and αCT motif together with the loop of FnIII-1 domain (blue) from the other protomer. Val3 of the insulin A chain (VA3, yellow) interacts with Asp707 in the IR (D707, red). **(D)** Insulin-interacting residues at IR site-1, shown in yellow. **(E)** Structural view of insulin (purple) bound to site-2 of the IR, engaging the FnIII-1 domain (blue). **(F)** Insulin-interacting residues at IR site-2, shown in purple. **(G)** AlphaFold-predicted structure of the IR bound to two RF-409 molecules. The protomers are shown in green and blue. RF-409 is shown in gray, with site-1 and site-2 components highlighted in yellow and purple, respectively. **(H)** Alphafold2 model of RF-409 binding to the IR, illustrating a non-canonical binding mode distinct from insulin. The L1 domain (green) of one IR protomer and the FnIII-1 domain (blue) of the other protomer are engaged by RF-409 (gray), with site-1 binding residues highlighted in yellow and the site-2 binding residues in purple. **(I)** Dose–response analysis showing EC_50_ values for IR phosphorylation (pY IR) and downstream signaling (pAKT and pERK). Log-transformed EC_50_ values (LogEC_50_) derived from dose-response curves in differentiated 3T3-L1 adipocytes (Fig. S1A,B) are shown as mean ± SEM, with corresponding EC_50_ values indicated for reference. **(J)** Lipogenesis assay in primary rat hepatocytes comparing metabolic responses elicited by insulin and RF-409. Data are presented as mean ± SEM; one-way ANOVA; *n* = 5 independent experiments. **(K)** Inhibition of gluconeogenesis in primary mouse hepatocytes by insulin and RF-409. Data are presented as mean ± SEM; one-way ANOVA; *n* = 6 independent experiments.

Insulin activates IR by binding two distinct sites (site-1 and site-2)(*2, 42–46*). Because mature IR exists as a stable dimer with two site-1 and two site-2 binding surfaces, up to four insulin molecules can bind a single IR dimer, inducing a symmetric T-shaped active conformation(*43*) (Fig. 1B). At site-1, one insulin molecule binds the L1 domain of one IR protomer together with the αCT motif and the FnIII-1 loop of the other protomer, thereby crosslinking the two protomers (Fig. 1C,D). Another insulin molecule binds site-2 on IR using the face of insulin opposite to its site-1–binding interface (Fig. 1E,F), promoting receptor conformational changes required for optimal activation.

In contrast to insulin, RF-409 activates IR through a distinct mode of engagement. Two molecules of RF-409 bind a single IR dimer, with each RF-409 molecule simultaneously engaging site-1 on one IR protomer and site-2 on the other protomer (Fig. 1G,H). The rigidity between the two IR-binding components of RF-409 enforces this geometry and stabilizes a unique, extended T-shaped active conformation. This mode of receptor engagement is fundamentally distinct from insulin-mediated activation and provides a structural basis for bypassing defects associated with impaired insulin binding in pathogenic IR mutants.

To compare the potency and efficacy of RF-409 with insulin, differentiated 3T3-L1 adipocytes were treated with increasing doses of each ligand. RF-409 exhibited EC_50_ values comparable to, or modestly lower than, those of insulin for IR autophosphorylation and phosphorylation of the downstream kinases protein kinase B (AKT) and extracellular signal-regulated kinase (ERK) (Fig. 1I; Fig. S1A,B). Consistent with insulin action, RF-409 robustly promoted lipogenesis (Fig. 1J) and suppressed glucose production (Fig. 1K) in primary hepatocytes. Together, these data demonstrate that RF-409 effectively activates IR in cultured cells and recapitulates key metabolic actions of insulin in glucose and lipid metabolism.

### RF-409 elicits insulin-like signaling with altered receptor-proximal phosphoregulation and temporal dynamics

To compare signaling elicited by RF-409 and insulin, C2C12-IR cells(*41*) were treated with 100 nM of both ligands and analyzed by LC–MS-based phosphoproteomics at early (10 min) and later (60 min) time points(*47*) (Supplementary Table 1). This analysis identified > 29,000 class I phosphorylated sites (localization probability > 0.75) across 4,156 protein groups. Principal component analysis (PCA) showed that RF-409–and insulin-treated samples clustered together at each time point, with separation driven primarily by time rather than ligand (Fig. 2A). Consistently, hierarchical clustering revealed strong similarity between insulin and RF-409 at 10 min, with modest divergence at 60 min, while basal samples formed a distinct cluster (Fig. S2A)

**Fig. 2.**
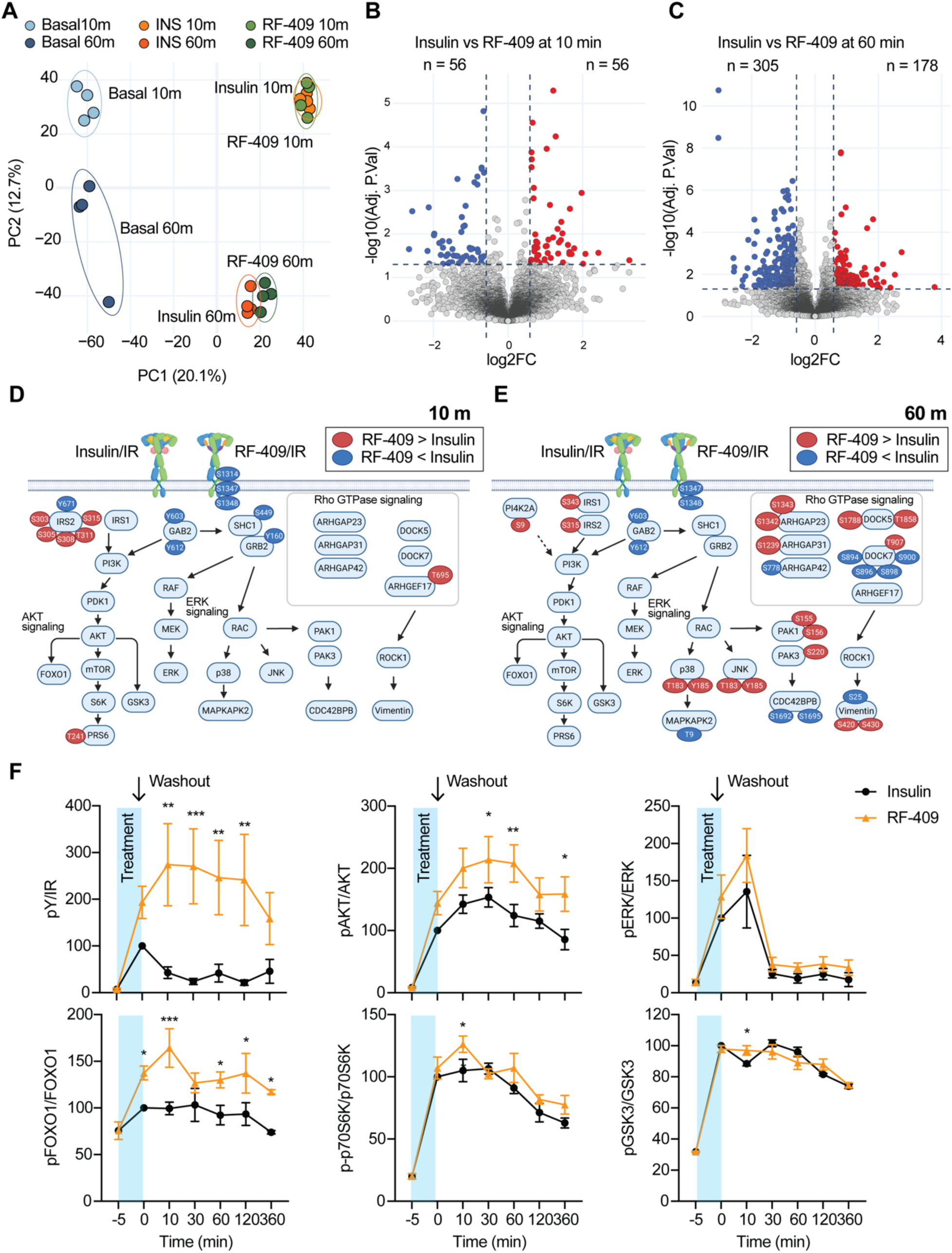
Phosphoproteomics reveals insulin-like signaling by RF-409. **(A)** Principal component analysis (PCA) of phosphoproteomic profiles from C2C12-IR cells treated with insulin or RF-409 at 10 min and 60 min (n = 4 per condition). **(B)** Volcano plot showing phosphosites differentially regulated between insulin- and RF-409-treated cells at 10 min. **(C)** Volcano plot showing phosphosites differentially regulated between insulin- and RF-409-treated cells at 60 min. **(D,E)** Integrated signaling map highlighting differentially regulated phosphosites between insulin and RF-409 at 10 min (D) and 60 min (E). Only significantly altered sites are shown for clarity. Early signaling events are largely shared between ligands, whereas differences emerge at receptor-proximal and adaptor-associated phosphorylation. Sites shown in red indicate higher phosphorylation in RF-409–treated samples relative to insulin, whereas sites shown in blue indicate lower phosphorylation. Arrows denote known protein–protein interactions and phosphorylation or dephosphorylation events curated from databases (PhosphoSitePlus) and the literature. (**F)** Prolonged IR signaling by RF-409. C2C12-IR cells were fasted for 4 h and treated with insulin or RF-409 (10 nM) for 5 min, followed by ligand washout and incubation for the indicated time points. Representative images are shown in Fig. S1C. Quantification of immunoblot data is shown. Data are presented as mean ± SEM; *n* = at least 3 independent experiments; Two-way ANOVA.

Both ligands induced widespread phosphorylation changes, with > 3,000 sites regulated relative to basal conditions at 10 min (Fig. S2B,C). In contrast, direct comparison between RF-409 and insulin identified only 112 differentially regulated phosphosites at 10 min, indicating highly similar phosphoproteomic profiles (Fig. 2B). At 60 min, differences became more apparent, with an increased number of differentially regulated sites, although the overall magnitude of divergence remained modest (Fig. 2C).

To visualize these differences, we generated an integrated signaling map highlighting phosphorylation changes between insulin and RF-409 (Fig. 2D,E). For clarity, only differentially regulated phosphosites are shown. Although our analysis achieved deep coverage across the pathway, inclusion of all detected sites reduced visual interpretability and is therefore omitted. At early time point, RF-409 closely recapitulated canonical insulin signaling, with comparable IR autophosphorylation on tyrosine residues and downstream activation of AKT and ERK pathways, indicating that, despite its distinct binding mode, RF-409 initiates an insulin-like signaling.

Despite this overall similarity, differences emerged in receptor-proximal phosphoregulation (Fig. 2D,E). Insulin robustly increased phosphorylation of multiple serine residues within the IR C-terminal region, including S1314, S1347, and S1348 (numbering based on human IR-B including signal peptide). In contrast, RF-409 induced weaker phosphorylation at S1314 and did not elevate S1347 and S1348 to the same extent at either time point. These findings suggest that RF-409 attenuates insulin-associated C-terminal serine phosphorylation of IR.

We next examined proximal adaptor signaling (Fig. 2D,E). IRS1 tyrosine phosphorylation was comparable between conditions. IRS1 S343 phosphorylation decreased at early time points under both conditions but was relatively higher with RF-409 at later time points. In contrast, IRS2 phosphorylation showed distinct regulation. At 10 min, insulin induced higher IRS2 Y671 phosphorylation than RF-409. Insulin also rapidly reduced phosphorylation of an IRS2 serine/threonine cluster (S303, S305, S308, T311, and S315), whereas RF-409 decreased these sites more gradually. By 60 min, phosphorylation remained reduced under both conditions but with distinct kinetics, indicating altered temporal control of adaptor phosphoregulation. Notably, S303 has been implicated in feedback regulation of IRS2 signaling(*48*), suggesting that RF-409 modifies proximal feedback timing. In addition, GAB2 phosphorylation at Y603 and S612 increased at early time points in response to both ligands, although the magnitude of induction was lower with RF-409. These results support differential regulation of adaptor-associated signaling downstream of IR.

At later time points, RF-409 was associated with differential regulation of cytoskeletal and trafficking pathways, including Rho GTPase and PAK signaling (Fig. 2E). Together, these results indicate that RF-409 largely recapitulates insulin signaling, with modest differences emerging over time in proximal phosphoregulation and feedback kinetics.

To determine whether the temporal differences observed by phosphoproteomics reflect altered signal duration, we examined the persistence of IR signaling following transient ligand exposure (Fig. 2F; Fig. S1C). C2C12-IR cells were treated with insulin or RF-409 for 5 min, washed extensively, and collected at the indicated time points. RF-409 sustained IR autophosphorylation for a longer duration than insulin. AKT phosphorylation was similarly prolonged after washout, with higher pAKT levels in RF-409–treated cells, whereas ERK phosphorylation showed limited persistence. Consistent with sustained AKT activity, downstream targets including FOXO1, GSK3β, and S6K remained phosphorylated, with modestly higher levels in RF-409–treated cells. These findings indicate that RF-409 prolongs IR signaling following transient stimulation, consistent with phosphoproteomic analyses, while maintaining broadly similar signaling outputs to insulin.

### Integrated pharmacokinetic and pharmacodynamic analysis of RF-409 in vivo

To characterize the in vivo pharmacokinetic (PK) and pharmacodynamic (PD) properties of RF-409, we performed integrated PK–PD analyses following systemic administration. For PD studies, insulin or RF-409 was administered subcutaneously, and insulin-responsive tissues, including liver, skeletal muscle, and epididymal white adipose tissue (eWAT), were collected for signaling analysis (Fig. 3A). IR activation and downstream pathway engagement were assessed by immunoblotting for IR autophosphorylation and phosphorylation of AKT and ERK.

**Fig. 3.**
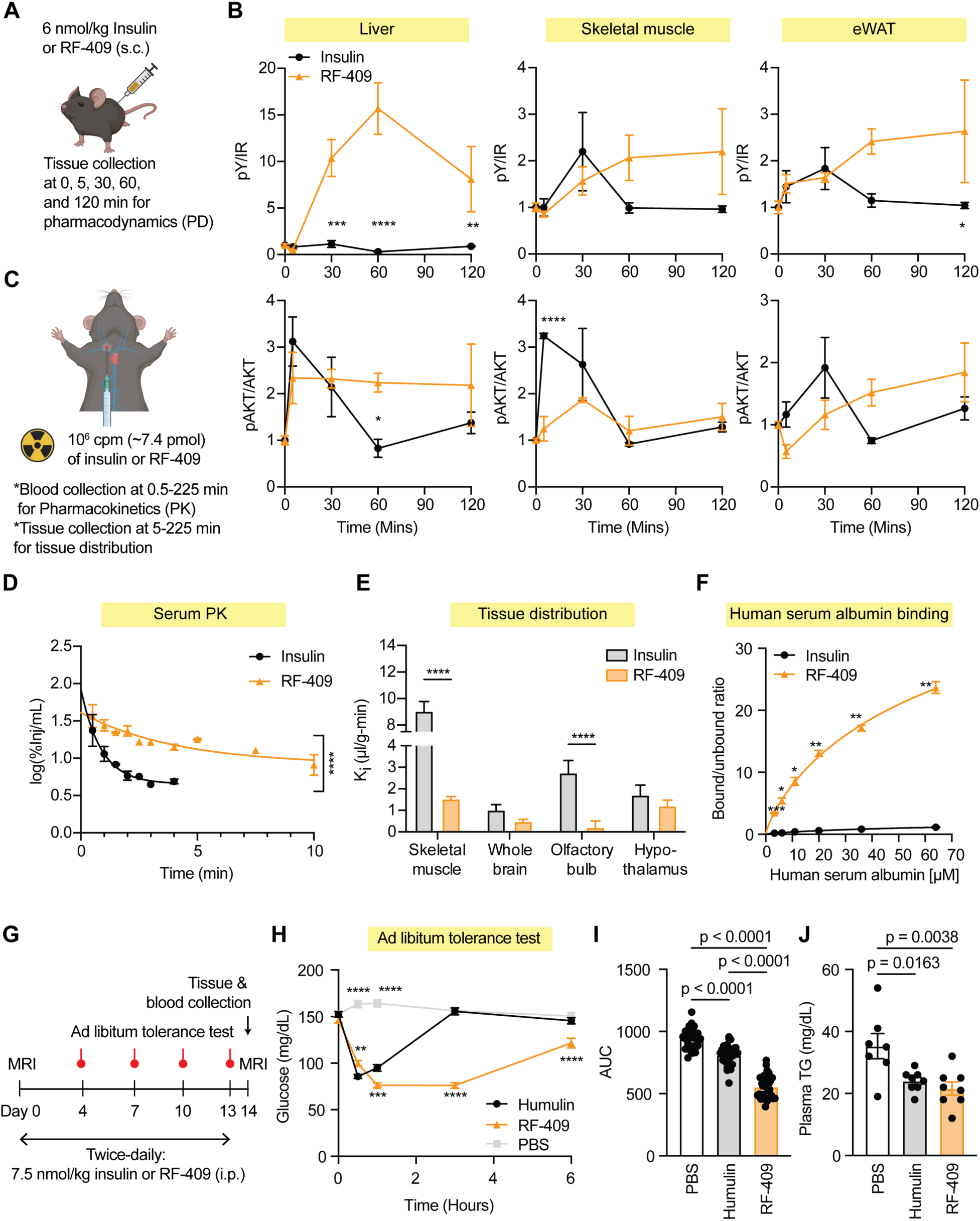
Pharmacokinetics and pharmacodynamics of RF-409. **(A)** Experimental scheme for pharmacodynamic studies shown in (B) and Fig. S3A–C. Mice were injected subcutaneously (s.c.) with insulin or RF-409 (6 nmol/kg), and tissues were collected at the indicated time points for pharmacodynamic analysis (*n* = 3 mice per time point). **(B)** Pharmacodynamic analysis of IR autophosphorylation and AKT phosphorylation in liver, skeletal muscle, and epididymal white adipose tissue (eWAT). For each sample, the ratio of phosphorylated to total protein was calculated and normalized to the reference value, defined as the average phosphorylated/total ratio of all samples collected at 0 min. Data are presented as mean ± SEM. *p < 0.05; **p < 0.01; ***p < 0.001; ****p < 0.0001 (Insulin vs. RF-409; Two-way ANOVA). **(C)** Experimental scheme for pharmacokinetic studies shown in (D), (E), and Fig. S3D–H. Mice were injected with radioisotope-labeled insulin or RF-409 (7.4 pmol) via the jugular vein. **(D)** Serum pharmacokinetics of insulin and RF-409. Data are presented as mean ± SEM; insulin, *n* = 16; RF-409, *n* = 28; two-way ANOVA; ****p < 0.0001. **(E)** Tissue distribution of insulin and RF-409. Data are presented as mean ± SEM. Insulin: skeletal muscle (*n* = 8), whole brain (*n* = 18), olfactory bulb (*n* = 10), hypothalamus (*n* = 17); RF-409: skeletal muscle (*n* = 20), whole brain (*n* = 28), olfactory bulb (*n* = 27), hypothalamus (*n* = 26). Two-way ANOVA; ****p < 0.0001. **(F)** Human serum albumin binding of insulin and RF-409. Data are presented as mean ± SEM; *n* = 3; two-way ANOVA; *p < 0.05, **p < 0.01; ***p < 0.001. **(G)** Experimental scheme for assessing the metabolic effects of RF-409 in healthy mice shown in (H–J) and Fig. S4. Male mice (2–3 months old) fed normal chow were injected intraperitoneally (i.p.) twice daily with the indicated doses of insulin or RF-409 for 14 days. Ad libitum tolerance tests were performed on the indicated days. Body composition was measured by MRI on day 0 and day 14. **(H)** Ad libitum tolerance test. Random-fed mice were injected with PBS, insulin (7.5 nmol/kg), or RF-409 (7.5 nmol/kg). Blood glucose levels were measured at the indicated time points after injection. Data represent the combined results of four independent ad libitum tolerance tests and are presented as mean ± SEM. Tolerance tests performed on individual treatment days are shown in Fig. S4. **p < 0.01; ***p < 0.001; ****p < 0.0001 (Humulin vs PBS or Humulin vs RF-409; two-way ANOVA). **(I)** Area under the curve (AUC) analysis of the ad libitum tolerance test shown in (H). **(J)** Fasting plasma triglyceride levels measured at the end of the treatment period.

In the liver, RF-409 induced significantly higher IR autophosphorylation compared with insulin and promoted prolonged AKT activation (Fig. 3B). ERK phosphorylation was minimal for both insulin and RF-409 under the dose and time conditions tested (Fig. S3A). In skeletal muscle, RF-409 elicited slower-onset but prolonged IR autophosphorylation relative to insulin, accompanied by delayed AKT activation that ultimately reached comparable levels (Fig. 3B). In contrast to liver, ERK phosphorylation was robustly induced in skeletal muscle by both insulin and RF-409 (Fig. S3B). In eWAT, RF-409 induced sustained IR autophosphorylation and showed a trend toward slower but prolonged AKT activation (Fig. 3B), whereas ERK phosphorylation was not significantly induced by either ligand under the conditions tested (Fig. S3C). These PD data demonstrate that RF-409 engages IR in vivo and produces tissue-specific signaling dynamics characterized by prolonged receptor activation and sustained AKT signaling, with ERK activation dependent on tissue context.

To define PK properties underlying these signaling profiles, isotope-labeled insulin or RF-409 was administered via jugular vein injection, and circulating radioactivity was quantified at multiple time points as a surrogate for ligand levels (Fig. 3C). Consistent with previous reports(*49, 50*), insulin exhibited rapid clearance, with an apparent early half-life of approximately 1 min based on regression over the 0.5–3 min sampling window and an estimated terminal half-life of ∼1.8 min derived from the 1.5–3 min interval (Fig. 3D; Fig. S3D). In contrast, RF-409 displayed markedly prolonged circulation, with an apparent half-life of ∼58 min based on regression over the 0.5–225 min interval and a terminal elimination half-life of ∼556 min (∼9.3 h) derived from ≥60 min time points. Because insulin was not sampled at later time points, its terminal elimination phase could not be fully resolved, and the calculated values likely underestimate its true terminal half-life; nonetheless, the difference in circulation time between insulin and RF-409 remains substantial.

To examine tissue distribution of RF-409, we focused on skeletal muscle where ligand delivery is constrained by endothelial transport(*51*), in contrast to the fenestrated liver sinusoid(*52*). RF-409 transport rate into skeletal muscle was significantly reduced relative to insulin (Fig. 3E; Fig. S3E). Given that RF-409 (∼16 kDa) is substantially larger than insulin (∼5.8 kDa), these differences in tissue distribution likely reflect, at least in part, size-dependent transport and tissue accessibility. Notably, despite reduced skeletal muscle transport, RF-409 elicited sustained IR signaling in this tissue (Fig. 3B), consistent with its prolonged systemic availability.

Given the distinct transport barriers governing brain entry, we next examined RF-409 distribution in the central nervous system. In the brain, RF-409 delivery to the whole brain was modestly reduced compared with insulin and was significantly lower in the olfactory bulb, a region exhibiting high rates of insulin transport(*53*), whereas hypothalamic levels were comparable between the two ligands (Fig. 3E; Fig. S3F-H). Notably, the hypothalamus contains a unique barrier transport system for insulin compared to the whole brain(*54*).

Albumin binding is a well-established mechanism for extending the half-life of peptide and protein therapeutics(*55–59*). To investigate whether the prolonged circulation of RF-409 reflects enhanced serum protein interactions, we performed human serum albumin (HSA) binding assays with insulin and RF-409 (Fig. 3F). RF-409 exhibited significantly higher binding affinity to HSA compared with insulin, consistent with increased plasma retention. These data suggest that enhanced albumin binding contributes to the extended systemic exposure of RF-409 observed in vivo. Together, these integrated PK–PD analyses indicate that RF-409 exhibits markedly prolonged circulation relative to insulin and produces sustained IR signaling across metabolic tissues in vivo.

### RF-409 produces sustained glucose lowering in healthy mice

RF-409 lowers glucose levels in both healthy and high-fat diet (HFD)–induced obese mouse models(*41*). To assess whether glucose-lowering activity is maintained during repeated administration, healthy mice were administered RF-409 twice daily for 14 days, with insulin and PBS groups treated under the same administration schedule (Fig. 3G). Ad libitum tolerance tests were performed on days 4, 7, 10, and 13 using the respective treatments (PBS, insulin or RF-409) to monitor glucose responses over the course of treatment. RF-409 produced a gradual reduction in glucose levels that was sustained over time (Fig. 3H,I; Fig. S4A-D). RF-409–treated mice exhibited robust glucose lowering at each time point, with no evidence of diminished activity during repeated administration.

Body composition was assessed by MRI prior to the first injection and again on day 14. No significant differences were observed in body weight, fat mass, or lean mass among PBS-, insulin-, and RF-409–treated groups (Fig. S4E-G), indicating that repeated RF-409 administration did not alter overall body composition in healthy mice.

Plasma chemistry analyses were performed at the end of treatment to assess metabolic and hepatic parameters. Both insulin- and RF-409–treated mice exhibited reduced plasma triglyceride levels compared with PBS controls (Fig. 3J), consistent with insulin action. Circulating albumin levels and liver enzyme markers, including alkaline phosphatase (ALP) and alanine aminotransferase (ALT), were comparable across all groups (Fig. S4H-J), and no evidence of overt inflammation or tissue pathology was observed. Collectively, these data show that RF-409 maintains glucose-lowering activity during repeated administration in healthy mice without detectable effects on body composition or liver function.

### Development of a severe insulin resistance mouse model

Severe insulin resistance syndromes caused by IR mutations represent a major clinical unmet need. A missense mutation substituting Asp707 with Ala in the IR has been identified in patients with Donohue syndrome(*60*). Asp707 is located within the αCT motif of IR (Fig. 1B, 1C, 4A), a structural element that is essential for insulin binding at site-1 and stabilization of the active receptor conformation(*43, 61*). Supporting this structural role, fibroblasts derived from the patient showed no detectable insulin binding(*60*), and insulin failed to activate the IR D707A mutant(*36, 40, 41, 43*). Importantly, the D707A mutation does not impair receptor processing or expression, indicating that the associated phenotypes arise primarily from defective insulin binding rather than reduced receptor abundance.

**Fig. 4.**
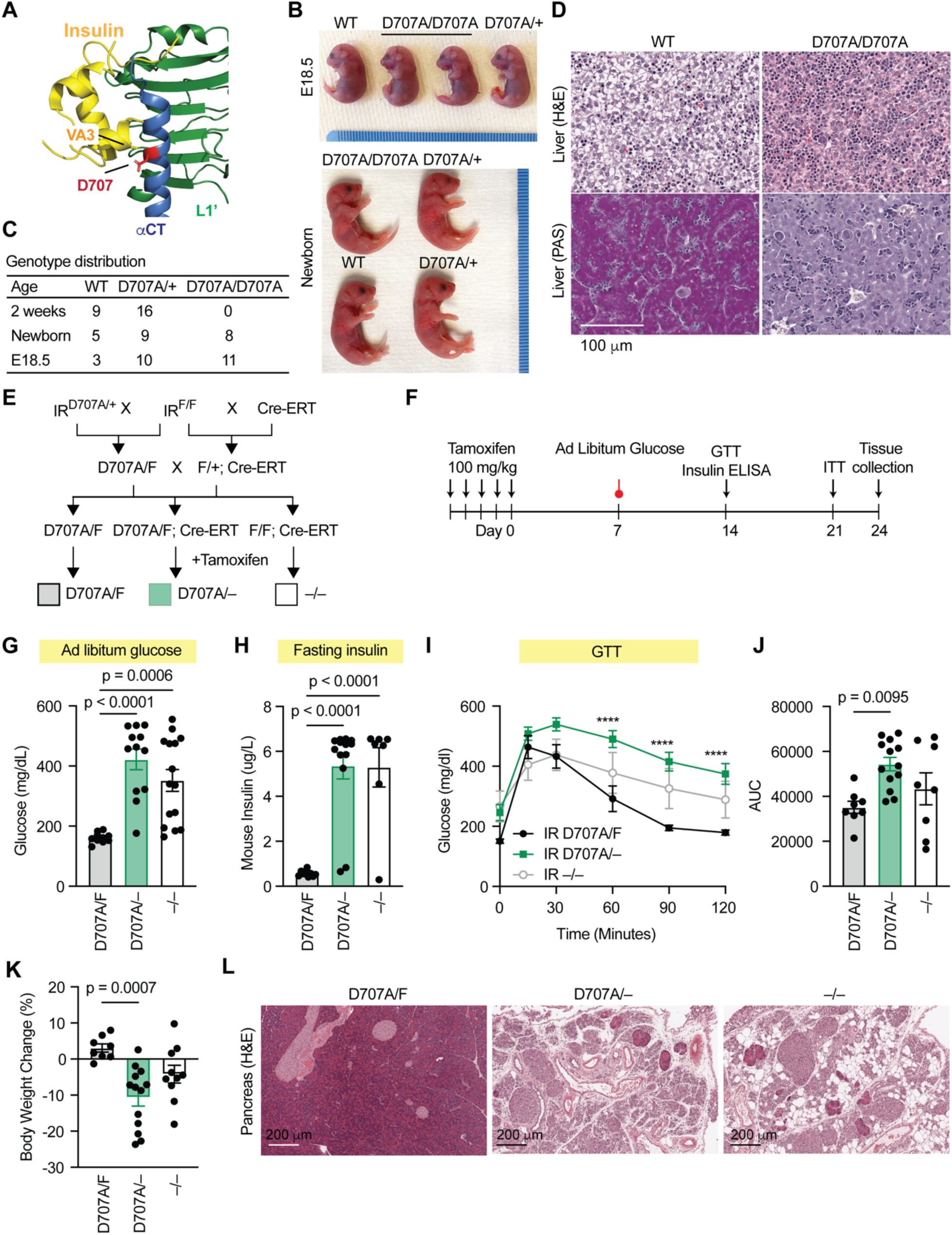
IR D707A mice exhibit severe insulin resistance. **(A)** Structural view of insulin binding at site-1 of the insulin receptor (IR). Asp707 (red) within the αCT motif (blue) of the IR is a key residue mediating site-1 insulin interaction. **(B)** Representative images of WT, D707A/+, and D707A/D707A littermates at embryonic day 18.5 and at birth. **(C)** Genotype distribution of WT, D707A/+, and D707A/D707A mice at different ages. **(D)** Hematoxylin and Eosin (H&E) staining and periodic acid–Schiff (PAS) staining (magenta) of livers from WT and D707A/D707A newborn mice. **(E)** Breeding strategy used to generate conditional D707A/– mice. **(F)** Experimental scheme. Male and female mice (2–3 months old) were injected with tamoxifen for five consecutive days to delete the floxed WT IR allele and generate D707A/– mice. Data from male mice are shown in (G-L) and data from female mice are shown in Fig. S5A–D. **(G)** Ad libitum blood glucose levels. D707A/F, *n* = 10; D707A/–, *n* = 12; –/–, *n* = 15. Data are presented as mean ± SEM; one-way ANOVA. **(H)** Fasting insulin levels. D707A/F, *n* = 9; D707A/–, *n* = 13; –/–, *n* = 7. Data are presented as mean ± SEM; one-way ANOVA. **(I)** Glucose tolerance test (GTT). Data are presented as mean ± SEM; two-way ANOVA; *n* = 8 mice per group; ****p < 0.0001. **(J)** Area under the curve (AUC) analysis of the GTT shown in (I). Data are presented as mean ± SEM; one-way ANOVA. **(K)** Percent change in body weight between pre-tamoxifen injection and the final day of the experiment. D707A/F, *n* = 8; D707A/–, *n* = 13; –/–, *n* =10. Data are presented as mean ± SEM; one-way ANOVA. **(L)** Representative H&E staining of pancreatic tissue.

Because RF-409 does not engage the αCT motif of IR and activates IR through a distinct binding mechanism (Fig. 1G,H), we asked whether it could bypass this defect in vivo. The residue corresponding to human IR D707 is Asp709 in mice. We therefore introduced an alanine substitution at this position to generate IR D709A knock-in mice (hereafter referred to as IR D707A mice). IR D707A/D707A mice exhibited no overt growth retardation or gross developmental abnormalities at birth (Fig. 4B,C). However, IR D707A/D707A pups died within 24 hours of birth.

To investigate the basis of this neonatal lethality, we examined metabolic defects in IR D707A mice. IR D707A/+ mice were indistinguishable from wild-type (WT) littermates (Fig. 4B). In contrast, although overall liver architecture appeared grossly normal, hepatocytes from IR D707A/D707A embryos (E18.5) and newborns displayed markedly reduced cytoplasmic vacuolation compared with WT and heterozygous mice (Fig. 4D), suggestive of impaired hepatic glycogen storage. Consistent with this interpretation, periodic acid–Schiff (PAS) staining revealed a dramatic reduction in glycogen content in IR D707A/D707A livers (Fig. 4D). Given that hepatic glycogen serves as a critical energy source during the neonatal period(*62*), insufficient glycogen storage—resulting in inadequate energy to sustain respiration and to transition from placenta support to nursing—likely contributes to the observed neonatal lethality. These phenotypes closely resemble those reported in IR knockout mice(*63, 64*) and in patients with Donohue syndrome(*60, 65*), supporting the relevance of IR D707A/D707A mice as a model of severe insulin resistance.

To circumvent neonatal lethality and enable assessment of RF-409 in adult animals, we generated a conditional IR D707A mouse model. IR D707A/+ mice were crossed with IR floxed mice (IR F/F) and a tamoxifen-inducible Cre recombinase (Cre-ERT) line to generate IR D707A/F, IR D707A/F;Cre-ERT, and IR F/F;Cre-ERT mice (Fig. 4E). Tamoxifen administration resulted in the generation of IR D707A/F, IR D707A/–, and IR –/– genotypes (Fig. 4E,F). As expected, IR –/– mice developed severe hyperglycemia (Fig. 4G), hyperinsulinemia (Fig. 4H), and profound glucose intolerance (Fig. 4I,J). IR D707A/– mice exhibited similarly elevated glucose and insulin levels compared with IR D707A/F controls, reaching levels comparable to IR –/– mice, and displayed severe glucose intolerance (Fig. 4G-J).

Body weight was significantly reduced in IR D707A/– mice relative to IR D707A/F controls (Fig. 4K). Notably, eWAT was essentially absent in IR D707A/– mice. Similar metabolic and adipose phenotypes were observed in female mice (Fig. S5A-D), indicating that IR D707A/– animals develop severe insulin resistance characterized by dysregulated glucose homeostasis, hyperinsulinemia, and profound lipoatrophy in both sexes.

Insulin signaling in pancreatic acinar cells is known to protect against cytotoxic Ca^2+^ overload and necrotic cell death(*66, 67*). Pancreatic insufficiency has been reported in patients with Donohue syndrome(*68*). Accordingly, both IR D707A/– and IR –/– mice exhibited severe pancreatic atrophy accompanied by acinar cell loss, fibrosis, and inflammation (Fig. 4L), consistent with the role of insulin as an exocrine pancreatic growth factor. In addition, pancreatic islets were enlarged in IR D707A/– and IR –/– mice, consistent with chronic hyperinsulinemia. Exocrine pancreatic atrophy likely contributes to the weight loss and lipoatrophy observed in IR D707A/– mice.

Hepatic steatosis and hepatomegaly with hepatocellular abnormalities have been described in severe insulin resistance syndromes(*9, 69, 70*). Both IR D707A/– and IR –/– mice exhibited marked cytoplasmic vacuolization of hepatocytes (Fig. S5E). Hepatocytes were enlarged, pale, and densely vacuolated. Scattered apoptotic bodies and focal areas of hepatocyte dropout were present, whereas fibrosis was not observed. These findings establish IR D707A/– mice as a robust and physiologically relevant in vivo model of severe insulin resistance syndromes and provide a platform to evaluate the therapeutic potential of RF-409.

### RF-409 activates IR D707A and restores insulin receptor signaling in vivo

To determine whether RF-409 can bind and activate the insulin-binding–deficient IR D707A mutant in vivo, PBS, Humulin, or RF-409 was administered via the inferior vena cava, and insulin-responsive tissues, including liver, skeletal muscle, and eWAT, were collected for signaling analysis (Fig. 5A). Because untreated IR D707A/– mice rapidly develop severe insulin resistance and poor health, signaling studies were performed 1 week after the final tamoxifen injection, a time point chosen to balance animal viability with receptor deletion efficiency.

**Fig. 5.**
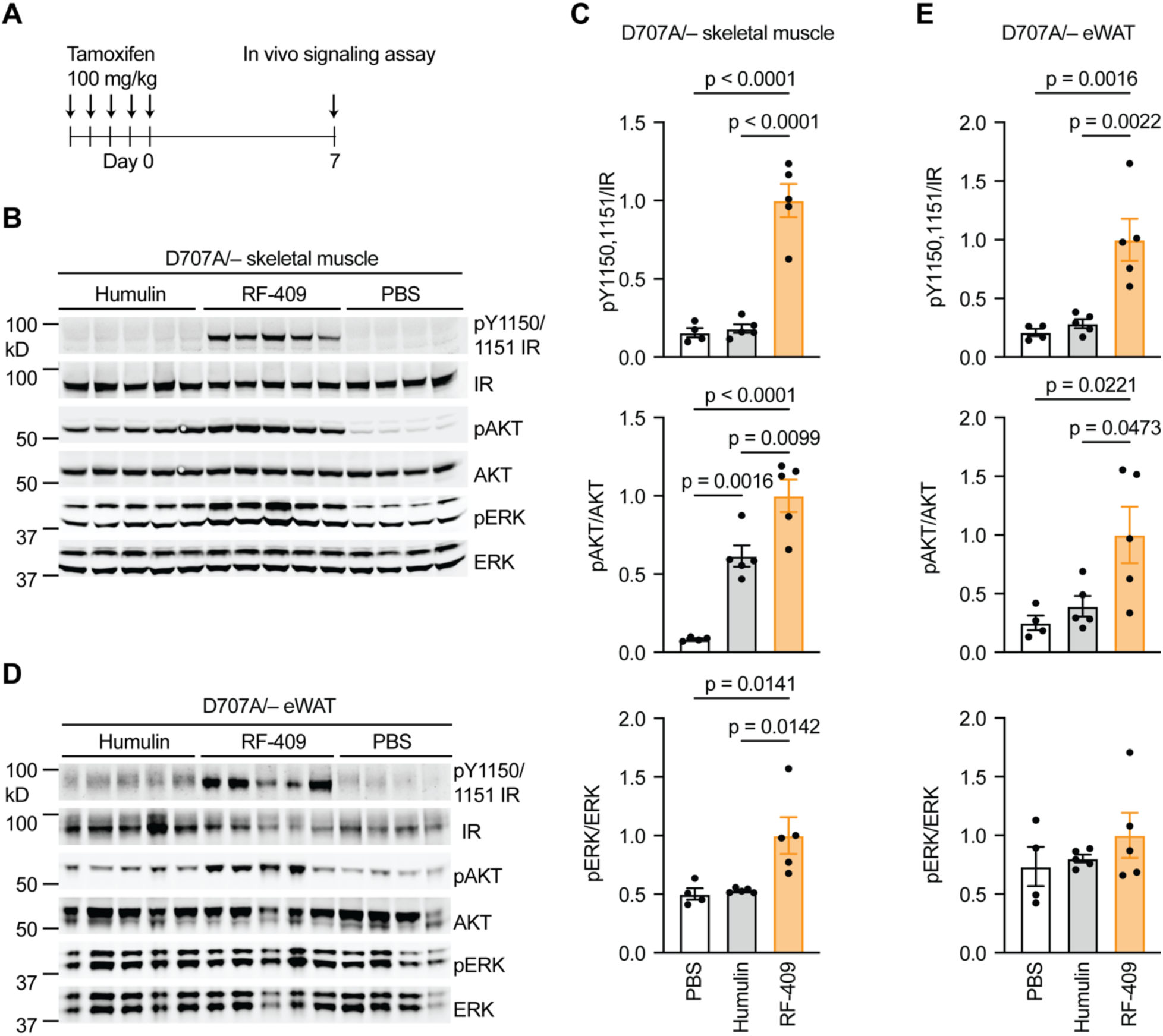
RF-409, but not insulin, activates IR D707A in vivo. **(A)** Experimental scheme. Female mice (2–3 months old) were injected with tamoxifen for five consecutive days to delete the floxed WT IR allele and generate D707A/– mice. Mice were fasted for 4 h and injected with PBS, 6 nmol/mouse Humulin or RF-409 via the inferior vena cava (IVC). Liver, skeletal muscle, and adipose tissue were collected at 3, 5, and 7 min after injection, respectively. **(B)** Representative immunoblot of IR autophosphorylation, AKT phosphorylation (pAKT), and ERK phosphorylation (pERK) in skeletal muscle from IR D707A/– mice. **(C)** Quantification of immunoblot data shown in (B). Data are presented as mean ± SEM; PBS, *n* = 4; Humulin, *n* = 5; RF-409, *n* = 5; one-way ANOVA. **(D)** Representative immunoblot of IR autophosphorylation, pAKT, and pERK in adipose tissue from IR D707A/– mice. **(E)** Quantification of immunoblot data shown in (D). Data are presented as mean ± SEM; PBS, *n* = 4; Humulin, *n* = 5; RF-409, *n* = 5; one-way ANOVA.

As expected, Humulin failed to activate IR D707A in peripheral tissues. In skeletal muscle, RF-409 markedly increased IR autophosphorylation as well as phosphorylation of AKT and ERK (Fig. 5B,C). Although insulin cannot directly activate IR D707A, the pAKT signal observed after Humulin treatment likely reflects residual WT IR signaling due to incomplete recombination and/or activation of IGF1R or IR/IGF1R hybrid receptors, rather than direct activation of IR D707A. In eWAT, RF-409 significantly increased IR autophosphorylation and pAKT (Fig. 5D,E), whereas pERK showed a modest increase that did not reach statistical significance. Humulin did not significantly increase pAKT or pERK in eWAT.

In the liver, tamoxifen-induced recombination did not achieve complete IR deletion (Fig. S6A,B). This likely reflects both the shortened interval between tamoxifen administration and tissue collection necessitated by the severe metabolic phenotype of IR D707A/– mice and mosaic Cre-ERT expression in hepatocytes reported for this Cre line(*71*). Consequently, both insulin and RF-409 increased IR autophosphorylation and pAKT in whole-liver lysates, with RF-409 inducing higher activation levels (Fig. S6C,D). pERK was not detectably increased under either condition.

To directly assess signaling through IR D707A under conditions of efficient receptor deletion, primary hepatocytes were isolated from IR D707A/F;Cre-ERT mice (Fig. S6E,F). Deletion of the WT IR allele was induced by treatment with 4-hydroxytamoxifen (4-OHT), resulting in more efficient IR knockout. Under these conditions, RF-409—but not insulin—robustly induced IR autophosphorylation, demonstrating direct activation of the insulin-binding–deficient IR D707A mutant. These data demonstrate that RF-409 directly activates the IR D707A mutant and restores downstream insulin signaling in vivo, supporting its ability to bypass defective insulin binding in a severe insulin resistance model.

### RF-409 controls glucose and lipid metabolism in a severe insulin resistance model

To test whether RF-409 regulates glucose levels in the severe insulin resistance model, we performed insulin tolerance assays using Humulin or RF-409. Humulin effectively reduced glucose levels in IR D707A/F mice but failed to do so in IR D707A/– mice (Fig. 6A,B; Fig. S7A,B). In contrast, RF-409 significantly lowered glucose levels in both IR D707A/F and IR D707A/– mice (Fig. 6C,D; Fig. S7C,D), consistent with activation of the insulin-binding–deficient receptor in vivo. RF-409 had minimal effects on glucose levels in IR –/– mice, indicating that its glucose-lowering activity requires IR expression.

**Fig. 6.**
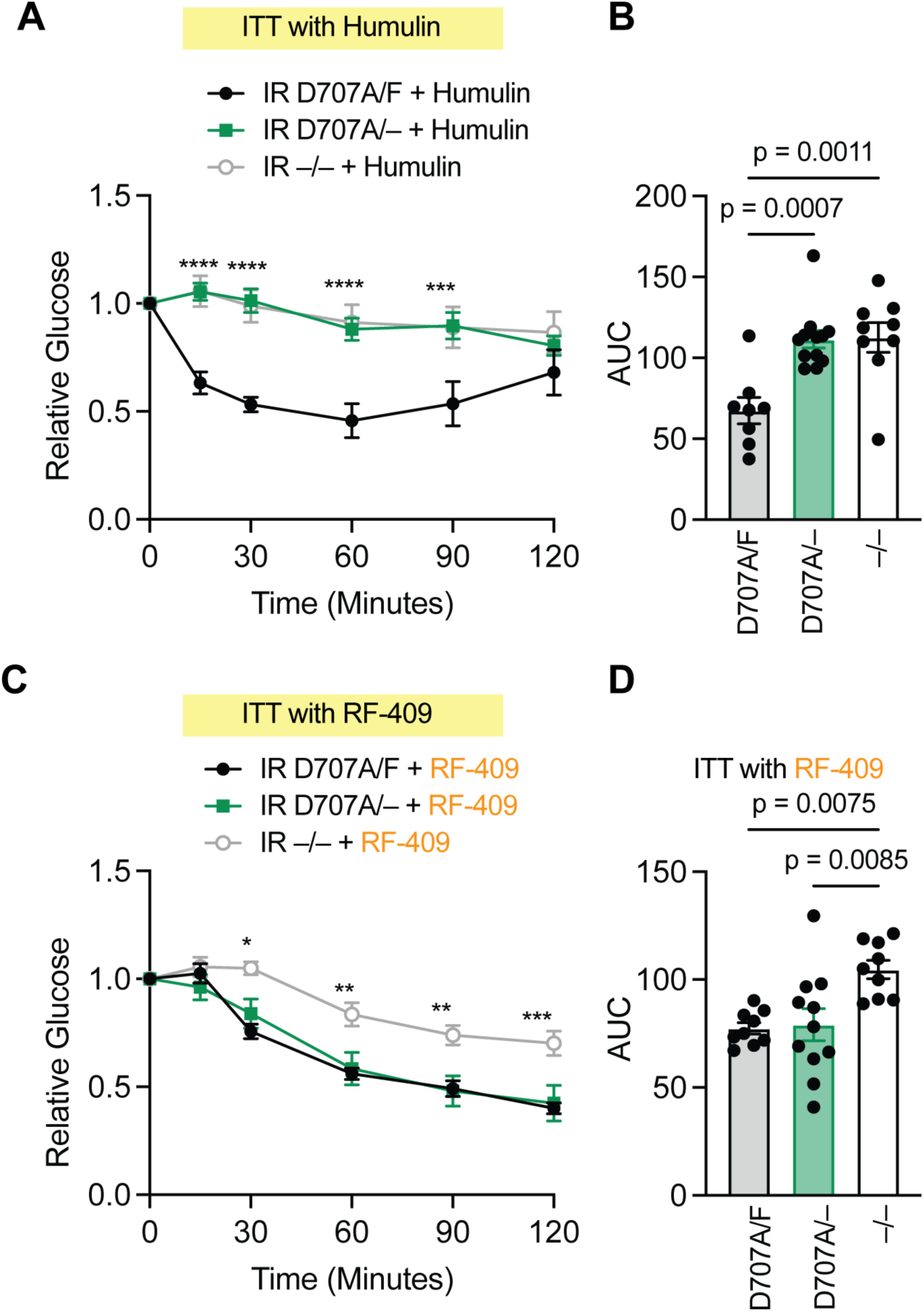
RF-409, but not insulin, lowers blood glucose levels in IR D707A mice. **(A)** Insulin tolerance test (ITT) following Humulin administration in male mice. IR D707A/F, *n* = 8; IR D707A/–, *n* = 12; IR –/–, *n* = 9. Data are presented as mean ± SEM; two-way ANOVA; ***p < 0.001, ****p < 0.0001 (IR D707A/F vs. IR D707A/–). **(B)** Area under the curve (AUC) analysis of the ITT shown in (a). Data are presented as mean ± SEM; one-way ANOVA. **(C)** Insulin tolerance test (ITT) following RF-409 administration in male mice. IR D707A/F, *n* = 9; IR D707A/–, *n* = 11; IR –/–, *n* = 9. Data are presented as mean ± SEM; two-way ANOVA; *p < 0.05, **p < 0.01, ***p < 0.001 (IR D707A/F vs. IR D707A/–). **(D)** Area under the curve (AUC) analysis of the ITT shown in (C). Data are presented as mean ± SEM; one-way ANOVA.

In female mice, but not male mice, Humulin produced a modest glucose-lowering effect in IR –/– animals (Fig. S7A,B), which may reflect incomplete deletion of both IR alleles following tamoxifen administration. Importantly, RF-409 significantly lowered glucose levels in IR D707A/– mice in both sexes but had no significant effect in IR –/– mice (Fig. 6C,D; Fig. S7C,D), demonstrating that RF-409 restores glucose regulation in mice harboring insulin-binding–deficient IR mutations, a model of severe insulin resistance.

We next examined the effects of chronic RF-409 treatment on systemic metabolic homeostasis in this severe insulin resistance model. Following five consecutive daily tamoxifen injections, osmotic pumps delivering PBS or RF-409 were implanted into IR D707A/– mice (Fig. 7A). PBS-treated IR D707A/– mice developed marked hyperglycemia (Fig. 7B,C) and hyperinsulinemia (Fig. 7D), and exhibited progressive body weight loss during the treatment period, accompanied by substantial reductions in both fat and lean mass (Fig. 7E-G). eWAT was nearly absent in PBS-treated IR D707A/– mice (Fig. 7H), consistent with severe lipoatrophy. Pancreatic atrophy and hepatic abnormalities were also evident in PBS–treated IR D707A/– mice (Fig. 7I-L).

**Fig. 7.**
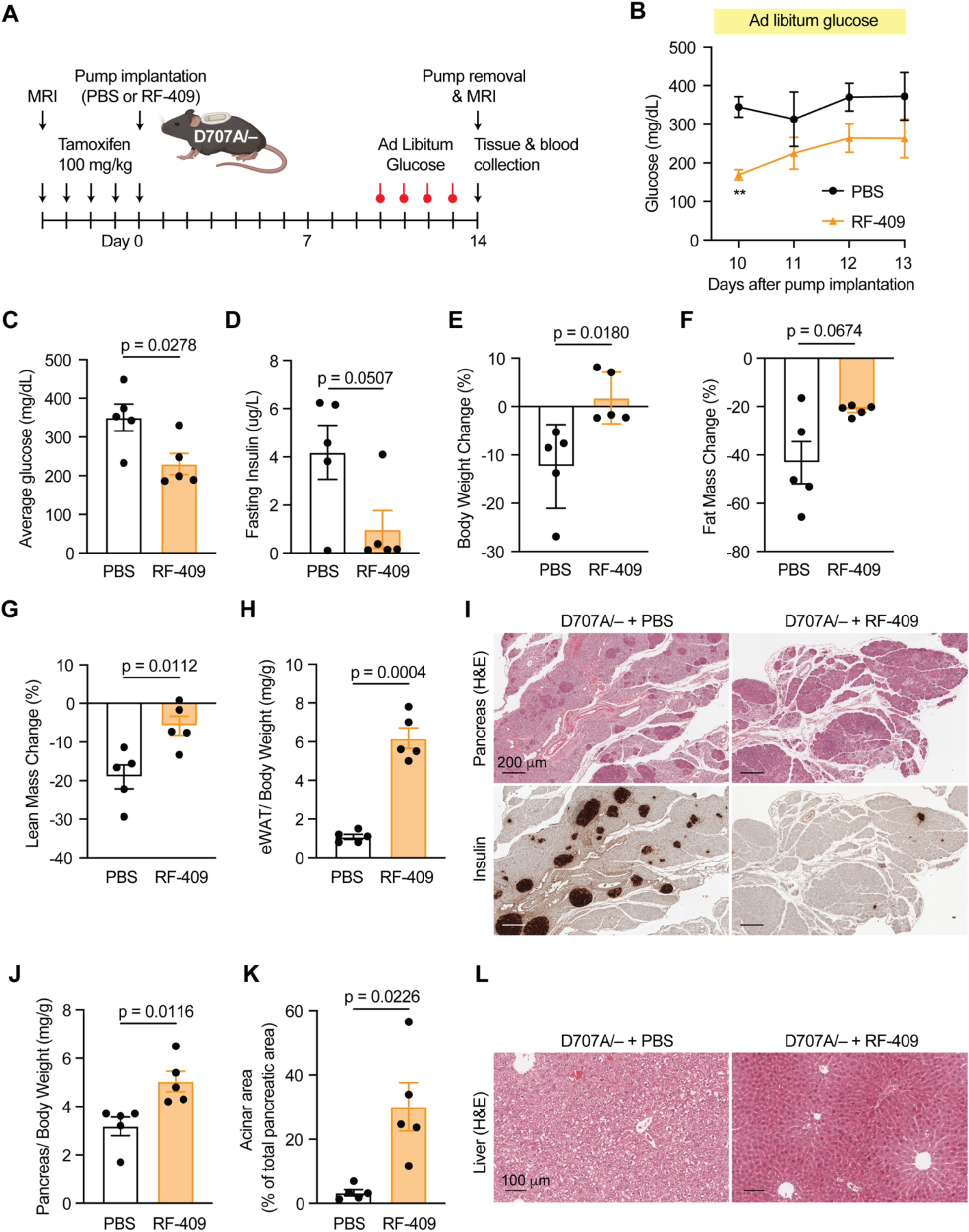
Chronic RF-409 treatment rescues severe insulin resistance phenotypes in IR D707A mice. **(A)** Experimental scheme for chronic RF-409 treatment. Male mice (3 months old) were injected with tamoxifen for five consecutive days to delete the floxed WT IR allele and generate IR D707A/– mice. Following the final tamoxifen injection, osmotic pumps releasing PBS or RF-409 were implanted. Body composition was assessed by MRI prior to tamoxifen injection and 14 days after the final tamoxifen injection. **(B)** Ad libitum blood glucose levels during the treatment period. Data are presented as mean ± SEM; *n* = 5 mice per treatment; two-way ANOVA; **p < 0.01. **(C)** Average ad libitum blood glucose levels calculated from (B). Data are presented as mean ± SEM; *n* = 5 mice per treatment; Welch’s t test. **(D)** Fasting insulin levels at the end of the treatment period. Data are presented as mean ± SEM; *n* = 5 mice per treatment; Welch’s t test. **(E)** Percent change in body weight over the treatment period. Data are presented as mean ± SEM; *n* = 5 mice per treatment; Welch’s t test. **(F)** Percent change in fat mass. Data are presented as mean ± SEM; *n* = 5 mice per treatment; Welch’s t test. **(G)** Percent change in lean mass. Data are presented as mean ± SEM; *n* = 5 mice per treatment; Welch’s t test. **(H)** Epididymal white adipose tissue (eWAT) weight normalized to body weight. Data are presented as mean ± SEM; *n* = 5 mice per treatment; Welch’s t test. **(I)** Representative H&E staining (top) and insulin immunostaining (bottom) of pancreatic tissue. **(J)** Pancreas weight normalized to body weight. Data are presented as mean ± SEM; *n* = 5 mice per treatment; Welch’s t test. **(K)** Acinar area quantification. Data are presented as mean ± SEM; *n* = 5 mice per treatment; Welch’s t test. **(L)** Representative H&E staining of liver tissue.

In contrast, chronic RF-409 treatment significantly reduced blood glucose levels (Fig. 7B,C) and normalized hyperinsulinemia (Fig. 7D). RF-409–treated mice maintained body weight, and both fat and lean mass were higher compared with PBS-treated controls (Fig. 7E-G). Moreover, lipoatrophy and pancreatic atrophy were markedly improved (Fig. 7H-K). Hepatocytes from RF-409–treated mice exhibited a marked reduction in cytoplasmic vacuolization compared with PBS-treated controls (Fig. 7L). Collectively, these data demonstrate that RF-409 restores insulin signaling and metabolic homeostasis in an insulin-binding–deficient severe insulin resistance mouse model and recapitulates key metabolic actions of insulin in vivo.

## DISCUSSION

Using structure-guided computational protein design, we developed a de novo IR agonist, RF-409, that activates IR through a noncanonical binding geometry. RF-409 recapitulates key metabolic actions of insulin in cultured cells and in vivo and exhibits prolonged systemic exposure associated with sustained, tissue-dependent signaling. In a patient-derived severe insulin resistance model harboring the IR D707A mutation, RF-409 bypasses insulin-binding deficiency, improves glucose control, and rescues systemic metabolic phenotypes following chronic administration.

Canonical insulin-mediated activation of IR involves multivalent engagement of site-1 and site-2 by multiple insulin molecules to stabilize an active receptor conformation. Although up to four insulin molecules can bind the receptor, distinct active conformations of IR may arise under sub-saturating ligand conditions, indicating that full ligand occupancy is not strictly required for signaling competence. Pathogenic mutations that disrupt these interactions, particularly at site-1, can uncouple ligand binding from receptor activation. Efforts to engineer insulin itself to overcome such defects are inherently limited, as the compact size and densely functional binding surface of insulin limit its engineerability for targeting mutant IRs. RF-409 overcomes this constraint by enforcing simultaneous engagement of two IR domains through two functional components connected by a rigid α-helix. This imposed geometry may mimic an activation-competent ligand-matured state, stabilizing an inter-protomer receptor conformation that enables activation of insulin-binding–deficient receptors such as IR D707A. Despite its distinct binding mode, RF-409 stabilizes an active IR conformation that resembles the insulin-induced relative orientation of the two membrane-proximal FnIII-3 domains, thereby promoting insulin-like downstream signaling.

RF-409 elicits insulin signaling that is initially comparable to insulin but diverges over time through altered receptor-proximal phosphoregulation and feedback kinetics. While early IR tyrosine phosphorylation and downstream AKT and MAPK activation were similar, RF-409 attenuated C-terminal serine phosphorylation of the receptor and altered adaptor phosphorylation, particularly within IRS2. One possible explanation is that RF-409 influences substrate engagement at the IR. Although IRS1 and IRS2 share PH/PTB-mediated docking, IRS2 can engage the receptor through additional determinants beyond the canonical interaction(*72*). This raises the possibility that ligand-dependent receptor states differentially support IRS1-versus IRS2-mediated signaling. Consistent with this, IRS1 tyrosine phosphorylation was unchanged, whereas early IRS2 Y671 phosphorylation was reduced with RF-409, suggesting altered substrate engagement kinetics. RF-409 also modified the temporal regulation of IRS2 serine/threonine phosphorylation, including sites linked to feedback control. Attenuation of receptor C-terminal serine phosphorylation, together with altered adaptor phosphoregulation, is consistent with reduced feedback-mediated desensitization and prolonged signaling output. These ligand-dependent differences at the receptor and adaptor levels may contribute to the sustained downstream signaling observed with RF-409.

A defining feature of RF-409 is its prolonged systemic exposure relative to insulin. Integrated pharmacokinetic analyses revealed circulation times on the order of hours rather than minutes, providing a pharmacokinetic basis for sustained tissue signaling and glucose-lowering activity. Enhanced binding to human serum albumin, together with increased molecular size, likely contributes to reduced clearance and extended exposure.

RF-409 transport into skeletal muscle was reduced; however, IR signaling remained sustained, suggesting that prolonged systemic availability and higher receptor affinity(*41*) may compensate for limited tissue penetration. RF-409 elicited the strongest IR activation in the liver, consistent with efficient ligand delivery in this highly perfused, fenestrated tissue. Because endogenous insulin is delivered to the liver via the portal circulation, whereas subcutaneous insulin therapy reverses this gradient, the liver-dominant signaling profile of RF-409 suggests partial restoration of a key feature of physiological insulin action—preferential hepatic engagement—even after systemic administration.

Since IR mutations were first identified in patients with severe insulin resistance in 1988(*73, 74*), these disorders have been recognized as devastating conditions associated with early morbidity and mortality. Despite multiple therapeutic strategies being explored, conventional treatments—including supraphysiologic insulin and recombinant human IGF-1—have failed to provide durable or mechanism-targeted benefit, and effective long-term therapies remain lacking. The ability to activate insulin-binding–deficient IR mutants using designer agonists therefore establishes a new translational avenue for congenital severe insulin resistance. RF-409 is not expected to be universally effective across all classes of IR mutations but is well suited to variants that retain receptor expression and intrinsic kinase activity while being defective in insulin binding or ligand-induced activation, as illustrated by the IR D707A mutation. Recent deep mutational scanning of the human IR ectodomain highlights the diversity of pathogenic mechanisms underlying insulin resistance and underscores the need for mutation-informed therapeutic strategies(*13*). Collectively, these observations support a precision-therapy framework that integrates structural prediction with functional screening to guide the use of engineered agonists such as RF-409.

RF-409 combines noncanonical receptor engagement with favorable pharmacokinetic properties and high IR selectivity without activation of the closely related IGF1R(*41*). These features are particularly relevant to pediatric patients, in whom genetic insulin resistance presents early in life and is refractory to existing therapeutic approaches. Beyond congenital syndromes, the prolonged circulation, receptor selectivity, and distinct engagement geometry of engineered IR agonists raise the possibility that such molecules could complement or, in specific contexts, provide an alternative to insulin therapy in more common forms of diabetes. Together, our findings highlight RF-409 as a targeted therapeutic strategy for severe insulin resistance syndromes and establish a broader framework for engineering receptor-selective insulin mimetics.

## MATERIALS AND METHODS

### Mouse strains and husbandry

All animal experiments described in this manuscript were approved by and conducted in accordance with the guidelines of the Institutional Animal Care and Use Committee (IACUC) at Columbia University. Pharmacokinetic (PK) studies were approved by and performed under the oversight of the VA Puget Sound Health Care System IACUC. Mice were maintained on a standard rodent chow diet (LabDiet #5053) and housed in a specific pathogen–free barrier facility under controlled environmental conditions (temperature, 68–70 °F; humidity, 30–70%) with a 12-hour light/dark cycle (lights on at 6:00 a.m., off at 6:00 p.m.).

### Generation of IR D707A/– mice and mouse husbandry

To generate IR D707A knock-in mice, a single-guide RNA (sgRNA) targeting the mouse *Insr* locus (5′-GTTGTGCAGGTAATCCTCGAAGG-3′), a single-stranded donor oligonucleotide containing the *Insr* p.D709A mutation (GAT→GCT) and a synonymous p.F707F mutation (TTC→TTT), and Cas9 protein were co-injected into fertilized C57BL/6N zygotes. Founder (F0) animals were identified by PCR amplification followed by Sanger sequencing and bred to wild-type (WT) mice to assess germline transmission and generate F1 offspring. Gene targeting and initial generation of IR D709A/+ mice (mouse numbering) were performed by Cyagen/Taconic. For consistency with human IR numbering, the mutation is referred to throughout this manuscript as IR D707A/+.

IR D707A/+ mice were backcrossed to a C57BL/6J background (The Jackson Laboratory, #000664) for at least three generations. *Insr* floxed mice (IR^F/F^; The Jackson Laboratory, #006955) were crossed with tamoxifen-inducible CAG-CreERT mice(*71*) (The Jackson Laboratory, #004682) to generate IR^F/F^;CreERT mice, which were subsequently backcrossed for nine generations onto the C57BL/6J background. Tamoxifen-inducible IR D707A/– mice were generated by crossing IR D707A/+ mice with IR^F/F^;CreERT mice.

### Genotyping

Genomic DNA was isolated from mouse tissue and PCR-amplified for genotyping. For D707A allele confirmation, PCR products were purified (Macherey-Nagel, #740609.250) and analyzed by Sanger sequencing. Cre genotyping was performed by PCR followed by agarose gel electrophoresis to identify Cre-positive animals. *Insr* floxed alleles were genotyped by PCR and resolved by agarose gel electrophoresis based on product size.

PCR primers used were as follows: D707A genotyping (D707A_F: CTCAGAAGCACAATCAGAGTGAGT; D707A_R: AGTGCACATGTGTGATCAGGTAG; D707A sequencing: AGTGCACATGTGTGATCAGGTAG). CreERT genotyping (Cre_F: AGGTTCGTTCACTCATGGA; Cre_R: TCGACCAGTTTAGTTACCC). IR floxed allele genotyping (IMR6765: GATGTGCACCCCATGTCTG; IMR6766: CTGAATAGCTGAGACCACAG).

### Cell lines

#### 3T3L1

3T3-L1 preadipocytes (ATCC CL-173) were cultured in high-glucose DMEM supplemented with 10% calf serum and 1% penicillin-streptomycin. Cells were seeded into 35 mm culture dishes and grown to full confluence prior to induction of differentiation. Differentiation was initiated by replacing the growth medium to differentiation medium consisting of high-glucose DMEM supplemented with 10% fetal bovine serum, 1% penicillin-streptomycin, 1.6 uM insulin (Sigma-Aldrich I2643), 1 uM dexamethasone (Sigma-Aldrich D4902), 0.5 mM 3-isobutyl-1-methylxanthine (Sigma-Aldrich D5879), and 10 mM rosiglitazone (Sigma-Aldrich R2408). After three days, the differentiation medium was replaced with maturation medium (High-glucose DMEM with 10% FBS, 1% penicillin-streptomycin, and 1.6 uM insulin) for an additional seven days. Prior to signaling assay, cells were weaned off insulin for 24 hours by culturing them in high-glucose DMEM with 10% FBS and 1% penicillin-streptomycin.

#### C2C12-IR

IGF1R knockout C2C12 cells expressing human IR-A(*41*) (C2C12-IR) were cultured in high-glucose DMEM supplemented with 10% (v/v) FBS, 2 mM L-glutamine, and 1% penicillin/streptomycin and maintained in monolayer culture at 37 °C and 5% CO2 incubator.

#### Cell line validation

Following passage of an aliquot of 3T3L1 for two weeks, a fresh batch of cells was thawed and propagated. There were no signs of mycoplasma contamination.

#### Lipogenesis assay

Primary hepatocytes from male rats were obtained from Thermo Fisher Scientific (RTCS10). Cells were thawed according to the manufacturer’s instructions and plated on collagen (Sigma, Cat. #C3867)-coated 35-mm culture dishes at a density of 5 × 10^5^ cells per dish. Hepatocytes were incubated in attachment medium (Williams’ E medium supplemented with 5% FBS, 10 nM insulin, 10 nM dexamethasone, and penicillin/streptomycin) for 5 hours, followed by overnight fasting in serum-free medium (DMEM low glucose supplemented with 0.5% BSA and penicillin/streptomycin).

After fasting, cells were pretreated for 2 hours with 100 nM insulin, RF-409, or vehicle control in serum-free medium. De novo lipogenesis was initiated by incubating cells for an additional 4 hours in serum-free medium containing the respective treatments and 0.6 µCi/mL [1,2-^14^C]-acetic acid sodium salt. Cells were washed three times with PBS to remove unincorporated tracer, and lipids were extracted twice using a 3:2 (v/v) hexane:isopropanol solution for 2 hours. The organic solvent was evaporated under nitrogen gas, and extracted lipids were resuspended in 2:1 (v/v) chloroform:methanol. Incorporated ^14^C radioactivity was quantified by liquid scintillation counting.

### Purification of RF-409

RF-409 was purified as previously described, with minor modifications(*41*). RF-409 was cloned with a N-terminal His6x tag with a TEV cleavage site and C-terminal FLAG tag in a pet29 based vector. RF-409 was expressed in BL21(DE3) *E. coli* (NEB C2527H) using autoinduction TBII medium (MP Biomedicals) supplemented with 50× 5052 and 20 mM MgSO_4_ under antibiotic selection. Cultures were grown at 37 °C for 6–8 h and subsequently shifted to 25 °C for overnight protein expression.

Cells were harvested by centrifugation at 4,000 × g and resuspended in lysis buffer (20 mM Tris, 300 mM NaCl, 5 mM imidazole, 0.7 % CHAPS, pH 8.0) supplemented with protease inhibitors (Thermo Fisher Scientific) and bovine pancreatic DNase I (Sigma-Aldrich). Cells were lysed by sonication, and lysates were clarified by centrifugation at 14,000 × g for 30 min at room temperature.

RF-409 was purified by immobilized metal affinity chromatography (IMAC). Clarified lysates were incubated with 6 mL nickel–nitrilotriacetic acid (Ni–NTA) agarose resin (Qiagen) for each gram of cell pellet harvested for 20–40 min at room temperature. The resin was washed with 5–10 column volumes of lysis buffer followed by 5–10 column volumes of wash buffer (20 mM Tris, 300 mM NaCl, 20 mM imidazole, pH 8.0). Bound protein was then eluted with elution buffer (20 mM Tris, 300 mM NaCl, 250 mM imidazole, pH 8.0).

Eluted fractions were incubated with MBP–TEV protease (Addgene plasmid) at a 1:20 molar ratio and dialyzed overnight at room temperature against PBS supplemented with 0.5 mM TCEP and 0.7% CHAPS to remove the His₆ tag and reduce endotoxin levels. The dialysis cassette was then transferred to PBS and dialyzed overnight to remove CHAPS and TCEP. The mixture was subsequently heated at 95 °C for 20 min and centrifuged at 14,000 × g for 20 min to precipitate and remove MBP–TEV. The purity of the purified RF-409 protein was assessed by SDS–PAGE and liquid chromatography–mass spectrometry. Purified protein was concentrated to 0.5–10 mg/mL and either stored at 4 °C for short-term use or flash-frozen in liquid nitrogen and stored at −80 °C.

### Protein modeling and visualization

The RF-409/IR structural model was generated by superposition RF-409 design model with cryo-EM structure of RF-405/IR (PDB:9DN6), as RF-409 and RF-405 are structural homologs (Figure 1G). RF-409/IR site-1 and site-2 are design models predicted by AlphaFold 2(*41*). Structural visualization and figure preparation were performed using PyMOL.

### Alexa Fluor 488 labeling of insulin and RF-409

Insulin (Sigma) or purified RF-409 were labeled with Alexa Fluor™ 488 NHS ester following the manufacturer’s instructions. Briefly, 0.5 mg of labeling dye was dissolved in 50 µl DMSO and was slowly added to 1 ml of protein (∼10 mg/ml) in 0.1 M sodium bicarbonate buffer (pH 8.3). The reaction was incubated for 1 h at room temperature with gentle rocking. Excess dye was removed by size-exclusion chromatography using a Superdex Increase 75 10/300 column (Cytiva) equilibrated with PBS.

### Differentiated 3T3L1 insulin receptor signaling assay

Insulin receptor (IR) signaling assays were performed as previously described, with minor modifications(*41*). Differentiated 3T3-L1 adipocytes were serum-starved prior to stimulation. For dose–response experiments, cells were treated with insulin (Sigma, I2643) or RF-409 at the indicated concentrations diluted in serum-free high-glucose DMEM for 10 minutes. Untreated control cells were incubated with serum-free high-glucose DMEM alone for 10 minutes.

Following treatment, cells were lysed on ice for 1 hour in lysis buffer B (50 mM HEPES, pH 7.4; 150 mM NaCl; 10% glycerol; 1% Triton X-100; 1 mM EDTA; 10 mM sodium fluoride; 2 mM sodium orthovanadate; 10 mM sodium pyrophosphate; 0.5 mM dithiothreitol; and 2 mM phenylmethylsulfonyl fluoride), supplemented with cOmplete Protease Inhibitor Cocktail (Roche), PhosSTOP (Roche), and Turbo nuclease (25 U/mL; Accelagen). Lysates were clarified by centrifugation at 18,213 × g for 20 minutes at 4 °C, and supernatants were mixed with SDS–PAGE sample loading buffer.

Proteins were resolved by SDS–PAGE and analyzed by immunoblotting. Primary antibodies used were anti-phospho-IR (Tyr1150/1151) (1:2000; 19H7; Cell Signaling Technology, #3024), anti-IR (1:500; CT3; Santa Cruz Biotechnology, sc-57342), anti-AKT (1:2000; 40D4; Cell Signaling Technology, #2920), anti-phospho-AKT (Ser473) (1:2000; D9E; Cell Signaling Technology, #4060), anti-ERK1/2 (1:2000; L34F12; Cell Signaling Technology, #4696), and anti-phospho-ERK1/2 (1:2000; 197G2; Cell Signaling Technology, #4377). For quantitative immunoblotting, DyLight 800–conjugated anti-rabbit IgG (#5151) and DyLight 680–conjugated anti-mouse IgG (#5470) secondary antibodies (Cell Signaling Technology) were used.

Membranes were scanned using the Odyssey Infrared Imaging System (LI-COR). Phosphorylation levels were normalized to total protein levels and expressed relative to the response induced by 100 nM insulin.

### Transient ligand washout assay for IR signaling persistence

C2C12-IR cells were serum-starved in serum-free DMEM for 4 hours. The cells were then treated with either 100 nM insulin or RF-409 for 5 minutes, followed by two extensive washes by PBS. After washing, the medium was replaced with fresh serum-free DMEM, and the cells were rapidly frozen at the designated time points at −80 °C. Untreated baseline control cells were incubated in serum-free DMEM for 5 minutes prior to collection.

Cells were lysed in RIPA Lysis and Extraction Buffer (Thermo Scientific, #89900) supplemented with Halt™ Protease and Phosphatase Inhibitor Cocktail, EDTA-free (Thermo Scientific, #78441), and Turbo nuclease (25 U/mL; Accelagen). Lysates were clarified by centrifugation at 18,213 × g for 20 minutes at 4 °C. The resulting supernatants were mixed with NuPAGE LDS Sample Buffer (Invitrogen, #NP0007) and 2-mercaptoethanol (10%; Sigma-Aldrich, #M6250).

Proteins were then resolved by SDS–PAGE and analyzed by immunoblotting. Primary antibodies used were anti-phospho-IR (Tyr1150/1151) (1:2000; 19H7; Cell Signaling Technology, #3024), anti-IR (1:500; CT3; Santa Cruz Biotechnology, sc-57342), anti-AKT (1:2000; 40D4; Cell Signaling Technology, #2920), anti-phospho-AKT (Ser473) (1:2000; D9E; Cell Signaling Technology, #4060), anti-ERK1/2 (1:2000; L34F12; Cell Signaling Technology, #4696), and anti-phospho-ERK1/2 (1:2000; 197G2; Cell Signaling Technology, #4377), anti-phospho-GSK3 beta (Ser9) (1:1000; D85E12; Cell Signaling Technology, #5558), anti-GSK3 beta (1:1000; 3D10; Cell Signaling Technology, #9832), anti-phospho-p70 S6 Kinase (Thr389) (1:1000; 1A5; Cell Signaling Technology, #9206), anti-p70 S6 Kinase (1:1000; 49D7; Cell Signaling Technology, #2708), anti-phospho-FOXO1 (Ser256) (1:1000; Cell Signaling Technology, #9461), anti-FOXO1 (1:1000; D7C1H; Cell Signaling Technology, #14952). For quantitative immunoblotting, DyLight 800–conjugated anti-rabbit IgG (#5151), DyLight 680–conjugated anti-mouse IgG (#5470), DyLight 800–conjugated anti-mouse IgG secondary antibodies (#5257), and DyLight 680–conjugated anti-rabbit IgG (#5366) (Cell Signaling Technology) were used. Membranes were scanned using the Odyssey Infrared Imaging System (LI-COR).

Phosphorylation levels were normalized to total protein levels and expressed relative to the response induced by insulin following a 5-minute treatment.

### Mouse hepatocyte isolation

Primary hepatocytes were isolated from 2-month-old male mice (C57BL/6J, Jackson #000664) with a standard two-step collagenase perfusion procedure as described earlier with some modifications(*40, 44, 75, 76*) . Briefly, following anesthesia, the inferior vena cava was cannulated, and the liver was perfused with Liver Perfusion Medium (Thermo Scientific, Cat. #17701038) using peristaltic pump (perfusion rate as 3 ml/min). After 1-2 sec upon appearance of white spots in the liver, we cut the portal vein with scissors to wash out blood, and then the liver was perfused with 30 ml of Liver Digest Medium (Thermo Scientific, Cat. #17703034). Dissected liver was gently washed with low glucose DMEM and transferred to sterile culture dish containing 15 ml Liver Digest Medium. The isolated liver cells were filtered through the 70 μm cell strainer into a 50 ml tube. After centrifuge at 50 g for 5 min at 4 °C, cells were washed with cold low glucose DMEM three times. Cells were resuspended with attached medium [Williams’ Medium E supplemented with 5% (v/v) FBS, 10 nM insulin, 10 nM dexamethasone, and 1% penicillin/streptomycin]. A million isolated primary hepatocytes were seeded in 35 mm collagen (Sigma, Cat. #C3867)-coated dishes. After 3 hours, the medium was changed to low-glucose DMEM medium supplemented with 10% (v/v) FBS and 1% penicillin/streptomycin overnight.

### Glucose production assay

Glucose production assay was performed previously described with slight modification(*77*). Briefly, hepatocytes were then serum starved with low-glucose DMEM supplemented with 1% penicillin/streptomycin overnight. After washing hepatocytes twice with PBS, the medium was replaced with glucose production medium (DMEM without glucose, glutamine, and phenol red (Thermo Scientific, Cat. # A1443001) supplemented with 20 mM calcium lactate, 2 mM sodium pyruvate, 1% (w/v) fatty-acid free BSA, and 1% penicillin/streptomycin). 1 mL glucose production medium was added to each cell culture dish along with following additions: (1) vehicle, (2) 100 uM cAMP and 1 uM dexamethasone (Dex/cAMP), (3) Dex/cAMP and 10 nM human insulin (I2643, Sigma), (4) Dex/cAMP and 10 nM of RF-409. After 6 h of incubation, medium was collected, and glucose concentration was measured using Glucose (GO) Assay Kit (Sigma, Cat. #GAGO20) and subsequently normalized to protein content.

### Cell culture and lysis for LC-MS phosphoproteomics

C2C12-IR cells were maintained below ∼60% confluency to prevent differentiation and seeded in 96-well plates at a density of 6,000 cells per well in 100 µL complete medium. Cells were allowed to adhere and proliferate for 22 hours. Serum starvation was performed by removing the medium, washing once with 100 µL phosphate-buffered saline (PBS), and replacing with 100 µL serum-free medium, followed by incubation at 37 °C for 6 hours.

Insulin and RF-409 were prepared in serum-free medium at final concentrations of 100 nM. Treatment was initiated by replacing the starvation medium with 100 µL of ligand-containing medium per well. To synchronize harvesting, the 60-minute treatment group was initiated first, followed by the 10-minute treatment group 50 minutes later, allowing all samples to be collected simultaneously. Each condition was analyzed in four replicates.

At the end of treatment, plates were immediately placed on dry ice, and cells were washed twice with 100 µL ice-cold Tris-buffered saline (1× TBS, pH 7.4). Cells were lysed by addition of 19 µL ice-cold lysis buffer (2% sodium deoxycholate (SDC), 50 mM triethylammonium bicarbonate (TEAB), 10 mM tris(2-carboxyethyl)phosphine (TCEP), and 40 mM chloroacetamide (CAA), pH 8.5) per well, followed by brief centrifugation.

Samples were incubated at 75 °C for 15 minutes with shaking at 1,000 rpm using a thermoshaker, followed by brief centrifugation. Lysates were then sonicated for 5 minutes using alternating 10-second pulse and 10-second pause cycles. After a final centrifugation, samples were snap-frozen on dry ice and stored until further processing.

### Phosphopeptide enrichment

µPhos phosphopeptide enrichment was performed as previously described(*78*), with minor modifications. Briefly, 1 µL of trypsin–LysC was added to each well at concentration of 25 ng/µL. 1:100 enzyme-to-protein ratio. Plates were sealed with two layers of aluminum foil and incubated at 37 °C with shaking at 1,500 rpm for 2 hours. Following digestion, plates were briefly centrifuged, and 20 µL of 100% 2-propanol was added to each well. Plates were incubated for 30 seconds at 1,500 rpm at room temperature, followed by addition of 40 µL µPhos enrichment buffer. Samples were transferred to 96-well deep-well plates (Eppendorf), and 5 µL of TiO₂ suspension (1 mg/µL) was added to each sample. Plates were incubated at 40 °C with shaking at 1,500 rpm for 7 minutes. Samples were centrifuged at 1,500 × g for 1 minute, and supernatants were removed using a multichannel pipette. Beads were washed six times with 200 µL µPhos washing buffer. Beads were then transferred to in-house packed C8 StageTips in two steps by adding 75 µL µPhos transfer buffer, followed by centrifugation at 700 × g for 8 minutes. Phosphopeptides were eluted by two sequential additions of 30 µL µPhos elution buffer, each followed by centrifugation at 400 × g for 4 minutes. Eluates were vacuum-dried at 45 °C for 30 minutes until the remaining volume was less than 10 µL. EvoTips were prepared by washing once with 20 µL buffer B (99.9% acetonitrile, 0.1% formic acid), activating in 2-propanol for 1 minute, and equilibrating with 20 µL buffer A. After equilibration, 100 µL of buffer A was added to EvoTips. Concentrated samples were reconstituted in 240 µL Evosep buffer A (0.1% formic acid in ddH₂O) and loaded onto pre-equilibrated EvoTips, containing 100 µL buffer A. All centrifugation steps for EvoTip loading were performed at 700 × g for 1 minute, except for sample loading, which was performed for 4 minutes.

### LC–MS analysis

Samples were analyzed using an Evosep One LC system (Evosep) coupled to an Orbitrap Astral mass spectrometer (Thermo Fisher Scientific). Peptides were eluted from EvoTips using a Whisper Zoom gradient with a throughput of 120 samples per day on an Aurora Rapid column (5 cm length, 75 µm inner diameter, packed with 1.7 µm C18 beads; IonOpticks). The column temperature was maintained at 60 °C using a column heater (IonOpticks). The Orbitrap Astral was equipped with an EASY-Spray ion source (Thermo Fisher Scientific) and a FAIMS Pro interface (compensation voltage −38 V; carrier gas flow 3.5 L/min; Thermo Fisher Scientific). An electrospray voltage of 1,900 V was applied for ionization, and the radio frequency level was set to 40. Full MS (MS1) spectra were acquired over an m/z range of 380–1,380 at a resolution of 240,000 (at m/z 200), with a normalized automatic gain control (AGC) target of 500% and a maximum injection time of 3 ms. Data-independent acquisition (DIA) MS/MS scans were performed using 100 variable isolation windows designed with pyDIAid software(*79*) (Supplementary Table 1), with a maximum injection time of 10 ms. Fragmentation was performed using high-energy collisional dissociation (HCD) with a normalized collision energy of 25%.

### Spectral search and data analysis

LC–MS raw files were processed using Spectronaut v20.2 (Biognosys) without an experimental spectral library (directDIA+ workflow). Data were searched against the UniProt mouse reference proteome (accessed August 2024), supplemented with the human insulin receptor sequence (P06213). Trypsin was specified as the protease with a maximum of two missed cleavages, and a minimum peptide length of seven amino acids was required. Precursor and fragment ion mass tolerances were set to dynamic at both MS1 and MS2 levels. False discovery rates (FDRs) were controlled at ≤1% at PSM, peptide and protein levels during the directDIA+ library generation step, and at precursor and protein Q-value levels (< 1%) during targeted extraction. For phosphoproteomics analysis, cysteine carbamidomethylation was set as a fixed modification, while protein N-terminal acetylation, methionine oxidation, and serine/threonine/tyrosine (STY) phosphorylation were set as variable modifications using the BGS Phospho PTM workflow with PTM localization mode enabled. Spectronaut localization probability threshold was set to 0 to retrieve all identified phosphopeptides, and downstream filtering at localization probability was applied in Python as described below. Quantification values were filtered based on q-values (default threshold of 0.01), and cross-run normalization was performed using the Automatic normalization setting.

### Bioinformatics data analysis

Bioinformatics analyses were performed in Python (v3.11). Phosphoproteomics data were processed using Spectronaut v20.2 and exported with the default BGS Factory Report schema, including the additional columns ‘EG.PrecursorID’, ‘PEP.PeptidePosition’, ‘EG.PTMAssayProbability’, ‘PG.Genes’, and ‘PG.ProteinGroups’. Exported data were processed using a custom Python implementation of the Perseus PeptideCollapse plugin(*80*). Phosphosites with localization probability <0.75 were excluded. Intensities were log-transformed, and phosphosites with an identification rate <70% across samples were removed. Remaining missing values were imputed using K-nearest neighbors (KNN) imputation implemented in scikit-learn (v1.6.1). Principal component analysis (PCA) was performed on the filtered and imputed dataset using scikit-learn. Differential expression analysis was conducted using the limma R package via the rpy2 Python interface. Empirical Bayes variance moderation (eBayes) was applied, and multiple testing correction was performed using the Benjamini–Hochberg method with a false discovery rate (FDR) cutoff of 5%. Phosphosites were considered significantly regulated at adjusted P < 0.05 and |log2FC| > 0.585. Sample-to-sample Pearson correlation was computed on log2-transformed phosphosite intensities across all 27,250 quantified sites, left after identification rate filtering and KNN imputation, and visualized as a hierarchically clustered heatmap (Ward’s method, Euclidean distance). Data visualization was performed using the Plotly Python library.

### Pharmacodynamics

Mice were fasted for 4 hours, beginning at 08:00 h. Following the fast, mice were injected subcutaneously with either Humulin or RF-409 (6 nmol/kg). At the indicated time points after injection, mice were anesthetized with isoflurane, and tissues including liver, quadriceps skeletal muscle, and epididymal white adipose tissue were harvested.

Collected tissues were homogenized at a ratio of 50 mg tissue per 1 mL lysis buffer (50 mM HEPES, 150 mM NaCl, 10% [v/v] glycerol, 1% [v/v] Triton X-100, 1 mM EDTA, 0.5 mM dithiothreitol, and 2 mM phenylmethylsulfonyl fluoride), supplemented with cOmplete Protease Inhibitor Cocktail (Roche), PhosSTOP (Sigma-Aldrich), and TurboNuclease (25 U/mL; Accelagen), using a Fisherbrand Bead Mill homogenizer. Lysates were incubated on ice for 1 hour and subsequently clarified by centrifugation at 20,817 × g for 20 minutes at 4 °C. Protein concentrations were determined using the Micro BCA Protein Assay Kit (Thermo Fisher Scientific). Equal amounts of protein (100 µg per lane) were analyzed by quantitative immunoblotting using the LI-COR Odyssey Infrared Imaging System (LI-COR, Lincoln, NE).

Primary antibodies used were as follows: anti–phospho-IR (Tyr1150/1151) (1:1000; 19H7; Cell Signaling Technology, #3024), anti-IR (1:500; CT-3; Santa Cruz Biotechnology), anti-AKT (1:1000; 40D4; Cell Signaling Technology, #2920), anti–phospho-AKT (Ser473) (1:1000; D9E; Cell Signaling Technology, #4060), anti-ERK1/2 (1:1000; L34F12; Cell Signaling Technology, #4696), and anti–phospho-ERK1/2 (1:1000; 197G2; Cell Signaling Technology, #4377).

### Radioactive labeling of proteins

Insulin, RF-409, and albumin were radioactively labeled as previously described(*81*). Briefly, human insulin (10 µg; Sigma-Aldrich, I2643) or RF-409 was radioiodinated with 1 mCi Na^125^I (PerkinElmer) using the chloramine-T method (Sigma-Aldrich). Bovine serum albumin (BSA; Sigma-Aldrich) was radiolabeled with ^99m^Tc (GE Healthcare). ^125^I-insulin, ^125^I-RF-409, and ^99m^Tc-albumin were purified using Sephadex G-10 columns (Sigma-Aldrich). Quality control steps were performed to verify radiochemical integrity, including acid precipitation with 30% trichloroacetic acid (TCA; Sigma-Aldrich) to confirm intact radiolabeled substrates.

### Pharmacokinetics

On the day of the terminal study, male and female CD-1 mice were anesthetized with urethane (40%; 0.15 mL intraperitoneal injection) to minimize pain and distress. Tracer amounts of **^125^**I-insulin (1 × 10^6^ cpm) or ^125^I-RF-409 (1 × 10^6^ cpm) were co-injected intravenously with ^99m^Tc-albumin (5 × 10^5^ cpm) in 1% BSA/lactated Ringer’s solution (total volume 0.2 mL). ^99m^Tc-albumin was included as a marker of vascular space(*82*). Tracers were allowed to circulate for 0.5–10 minutes following injection.

Extended pharmacokinetic studies were performed for RF-409 by injecting ^125^I-RF-409 and collecting samples over a time course of 5–225 minutes. Terminal arterial blood samples were collected from the left carotid artery at multiple time points (0.5–225 minutes). Mice were then decapitated, and whole brains were rapidly removed, with the olfactory bulb and hypothalamus separated and weighed. Gastrocnemius muscle was collected from the left hindlimb and weighed.

Arterial blood samples were centrifuged at 5,400 × g for 10 minutes, and serum was collected. Radioactivity in serum (50 µL), brain regions, and muscle samples were quantified using a gamma counter (Wizard^2^, PerkinElmer).

Plasma concentration–time data for insulin and RF-409 were analyzed using noncompartmental methods(*83*). The percent of the injected dose of ^125^I-insulin or ^125^I-RF-409 present in one ml of serum (%Inj/ml) was calculated and its log value plotted against time as the radioactivity measurements spanned several orders of magnitude. For each ligand, an apparent terminal elimination phase was identified visually as the final time points that exhibited a linear relationship between log_10_ (C) and time. Linear least-squares regression was then performed on log_10_ (C) versus time to obtain the elimination slope (m, in log_10_ units per minute). The elimination half-life (t_1/2_) was calculated using:

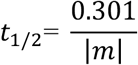

which is mathematically equivalent to 𝑡_1/2_ = 0.693/𝑘 when natural logarithms are used. All regressions and plots were generated using GraphPad Prism.

The tissue/serum (T/S) ratios were graphically displayed against their respective exposure times (Expt). Expt was calculated from the formula:

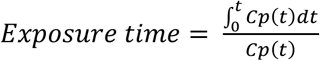

where *Cp* is the level of radioactivity (cpm) in serum at time (*t*). Expt corrects for the clearance of tracer from the blood. The influx of insulin and RF-409 was calculated by multiple-time regression analysis as described by Patlak, Blasberg, and Fenstermacher(*82, 84*):

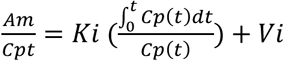

where *Am* is level of radioactivity (cpm) per g of brain tissue at time *t*, *Cpt* is the level of radioactivity (cpm) per ml arterial serum at time *t*, K*_i_* (μl/g-min) is the steady-state rate of unidirectional solute influx from blood to brain, and V*_i_* (μl/g) is the level of rapid and reversible binding for brain which usually is a combination of vascular space plus any brain endothelial cell receptor binding. The tissue/serum (T/S) ratios for insulin and RF-409 were corrected for vascular space by subtracting the corresponding ratio for albumin, yielding a delta T/S ratio. The linear portion of the relation between the delta T/S ratio versus Expt was used to calculate the K*_i_* (μl/g-min) with its standard error term, and the y-intercept determined as representation of the V*_i_* (μl/g)(*82*).

Regression analyses and statistical comparisons were performed using Prism 10 (GraphPad Software, San Diego, CA, USA). For pharmacokinetic studies, slopes (𝐾_i_), correlation coefficients (r), and y-intercepts (𝑉_i_) were compared statistically using Prism as previously described(*85*).

### Human serum albumin binding assay

Relative binding of insulin and RF-409 to human serum albumin (HSA) was assessed using the TRANSIL HSA Binding Kit (Sovicell) according to the manufacturer’s instructions. Briefly, Alexa Fluor 488–labeled insulin or RF-409 was incubated with increasing amounts of HSA-conjugated beads such that the final concentration of each ligand was 100 nM. Samples were mixed at 1,000 rpm for 15 minutes using a MixMate instrument (Eppendorf) and subsequently centrifuged at 750 × g for 10 minutes to separate bead-bound from unbound ligand. The supernatant was collected, and fluorescence intensity was measured to quantify unbound ligand. Binding curves were generated using the manufacturer-provided analysis software.

### Metabolic characterization of RF-409 in healthy mice

Two-month-old male C57BL/6J mice (The Jackson Laboratory, #000664) were acclimated for one week prior to the start of the experiment. Mice received twice-daily subcutaneous injections of PBS (vehicle), insulin (7.5 nmol/kg), or RF-409 (7.5 nmol/kg) for the duration of the study. Body composition was assessed by magnetic resonance imaging (EchoMRI^TM^) at baseline and at the experimental endpoint to determine changes in fat and lean mass.

Ad libitum insulin tolerance tests (ITTs) were performed on experimental days 4, 7, 10, and 13. For each ITT, mice were injected with PBS, insulin (7.5 nmol/kg), or RF-409 (7.5 nmol/kg), and blood glucose levels were measured from tail vein blood at 0, 0.5, 1, 3, and 6 hours post-injection using a glucometer (Contour Next). ITT measurements from the four test days were combined for analysis.

At the experimental endpoint, mice were fasted for 5 hours, and terminal blood was collected by cardiac puncture under anesthesia for analysis of circulating metabolites. Blood was dispensed into heparin-treated tubes, centrifuged at 2,000 × g for 15 minutes, and plasma was isolated for blood chemistry analysis performed by Diagnostic Lab Services at Columbia University. Parameters measured included albumin, alkaline phosphatase (ALP), alanine aminotransferase (ALT), and triglycerides (TG).

### Tissue Histology

Mouse tissues were fixed in 10% neutral buffered formalin (NBF) for hematoxylin and eosin (H&E) staining or in 1% periodic acid prepared in 10% NBF for periodic acid–Schiff (PAS) staining. Tissue processing, embedding, sectioning, and staining were performed by the Molecular Pathology Core at Columbia University. Stained slides were scanned using a Leica SCN400 slide scanner. Pancreatic acinar area was quantified from histological sections using QuPath image analysis software.

### Immunohistochemistry

Tissues were fixed in 10% neutral buffered formalin and embedded in paraffin blocks by the Diabetes Research Core at Columbia University. Sections were deparaffinized, and heat-induced epitope retrieval was performed using 10 mM sodium citrate buffer (pH 6.0). A primary antibody against insulin (1:500; C27C9; Cell Signaling Technology, #3014) was used to label pancreatic β cells and islets. Immunohistochemistry was performed using the ImmPRESS HRP Goat Anti-Rabbit IgG Polymer Detection Kit (Vector Laboratories) according to the manufacturer’s instructions. Hematoxylin was used as a counterstain. Slides were scanned using a Leica SCN400 scanner by the Molecular Pathology Core at Columbia University.

### In vivo and in vitro Tamoxifen induction

For in vivo induction, mice were administered tamoxifen (100 mg/kg; Sigma-Aldrich, T28590) dissolved in corn oil by intraperitoneal injection once daily for five consecutive days. For in vitro induction, primary hepatocytes isolated from IR D707A/F;ERT-Cre mice were treated with 4-hydroxytamoxifen (EMD Millipore, 508225) at a final concentration of 1 μM for 3 days.

### Glucose tolerance test

Mice were fasted for 2 hours prior to testing. A glucose solution (2 g/kg body weight) was administered by intraperitoneal injection. Blood glucose levels were measured from tail vein blood at 0 (baseline), 15, 30, 60, 90, and 120 minutes post-injection using a glucometer. Glucose tolerance was quantified by calculating the area under the curve (AUC) using the trapezoidal method.

### Insulin tolerance test

Mice were fasted for 2 hours prior to testing. A bolus of insulin or RF-409 was administered by intraperitoneal injection. Blood glucose levels were measured from tail vein blood at 0 (baseline), 15, 30, 60, 90, and 120 minutes post-injection using a glucometer. Insulin sensitivity was quantified by calculating the area under the curve (AUC) of the glucose response using the trapezoidal method.

### Insulin ELISA

Mice were fasted for 5 hours prior to blood collection from the tail vein. Blood samples were centrifuged at 2,000 × g for 15 minutes, and plasma was isolated. Plasma insulin levels were measured using a Mouse Insulin ELISA Kit (Mercodia) according to the manufacturer’s instructions.

### In vivo insulin signaling assay in severe insulin resistance model

Cre-mediated recombination was induced by intraperitoneal injection of tamoxifen (100 mg/kg; Sigma-Aldrich, T28590) dissolved in corn oil once daily for five consecutive days. Seven days after the final tamoxifen injection, mice were anesthetized, and a bolus of PBS, insulin, or RF-409 (6 nmol/mouse) was administered via the inferior vena cava.

Liver, epididymal white adipose tissue, and skeletal muscle were harvested at 3, 5, and 7 minutes post-injection, respectively. Tissues were lysed in RIPA buffer (Thermo Scientific, 89900) supplemented with protease and phosphatase inhibitors (Thermo Scientific, 78442). Protein concentrations were determined using a BCA assay to ensure equal loading.

Protein lysates were analyzed by immunoblotting using the following primary antibodies: anti–phospho-IR (Tyr1150/1151) (1:1000; 19H7; Cell Signaling Technology, #3024), anti-IR (1:500; CT-3; Santa Cruz Biotechnology), anti-AKT (1:1000; 40D4; Cell Signaling Technology, #2920), anti–phospho-AKT (Ser473) (1:1000; D9E; Cell Signaling Technology, #4060), anti-ERK1/2 (1:1000; L34F12; Cell Signaling Technology, #4696), and anti–phospho-ERK1/2 (1:1000; 197G2; Cell Signaling Technology, #4377).

### Metabolic characterization of chronic RF-409 treatment in insulin resistant model

Baseline body weight, fat mass, and lean mass were assessed by magnetic resonance imaging prior to tamoxifen induction. Mice then received five consecutive daily intraperitoneal tamoxifen injections and were anesthetized with 3% isoflurane for subcutaneous implantation of an osmotic pump (Alzet 1002) delivering either PBS or 1.2 nmol/day RF-409. Ad libitum blood glucose levels were measured at 10:00 AM daily during the final 4 days of the treatment period. Fourteen days after pump implantation, body weight and composition were reassessed. Mice were subsequently fasted for 5 hours. Blood was collected by cardiac puncture under anesthesia and was then transferred to heparin-treated tubes. Plasma was obtained by centrifuging whole blood at 2000 x g for 15 minutes. Liver, pancreas, and eWAT were carefully dissected and weighed. Liver and pancreas tissues were fixed in 10% neutral buffered formalin for subsequent histological analysis.

### Statistical analysis

Graph generation and statistical analyses were performed using Prism 10 (GraphPad Software). Data are presented as mean ± SD or mean ± SEM, as indicated in the figure legends. Two-tailed unpaired Student’s *t* tests were used for pairwise comparisons. One-way or two-way ANOVA was performed as appropriate, followed by Tukey’s multiple-comparisons test. Sample sizes were based on the maximum number of mice available for each experiment. No formal power analysis was performed to predetermine sample size. Randomization and blinding were not performed. Data were analyzed after completion of data collection for each experiment.

## Acknowledgements

This work was supported in part by grants from the National Institutes of Health (R35GM142937 and R21TR006115 to E.C.; R01DK58282 to D.A.), the Irma T. Hirschl Award (to E.C.), and the Howard Hughes Medical Institute (to D.B.). A.H. was supported by an NIH F31 Predoctoral Fellowship (F31DK141231). M.G. was supported by an NIH MSTP T32 (T32GM145440). E.R. was supported by the VA Puget Sound Research and Development Service. This research also used resources supported by NIH funding to the Columbia Diabetes Research Center (P30DK063608) and the Columbia Digestive and Liver Diseases Research Center (P30DK132710). We thank Ana M. Felte Castro and Thomas Kolar for assistance with mouse maintenance and tissue processing, and Chuchun Liz Chang for sharing equipment for the lipogenesis assay. We are grateful to members of the Accili and Choi laboratories for insightful discussions.

## Author Contributions

E.C. and D.A. designed the research. X.W. and D.B. designed RF-409. X.W., M.G., and J.Y. purified RF-409. M.N. and C.P. performed signaling assays in cultured cells. M.N. maintained the mouse colonies. A.H. performed in vitro lipogenesis and glucose production assays. A.H., D.A., and E.C. characterized RF-409 function in mice and analyzed the animal experiments. M.G., E.R., and G.R. performed PK and PD analyses. E.R. and R.W. performed and analyzed tissue distribution of RF-409 in mice. D.O., E.Z., and M.M. performed and analyzed phosphoproteomics. Q.F. provided consultation on protein purification. E.C. wrote the initial manuscript, and all authors contributed to editing and discussion of the manuscript.

## Competing interests

X.W., E.C., S.C., and D.B. are co-inventors on a provisional patent (IP: 50206.01US1) filed by the University of Washington covering RF-409 and its uses described in this manuscript. All other authors declare no competing interests.

---

**Fig. S1.**
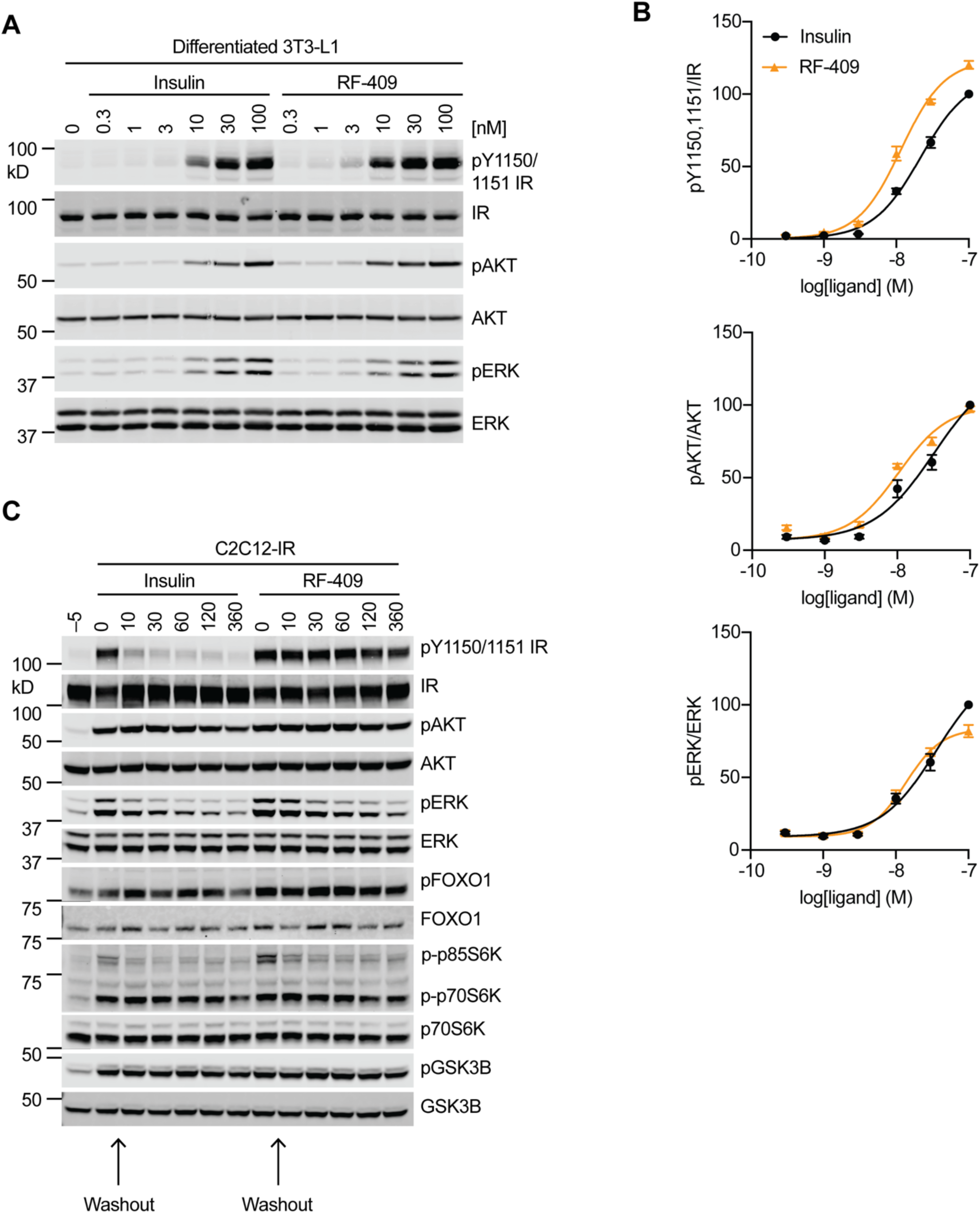
RF-409 potently activates and prolongs IR signaling. **(A)** Representative immunoblot of differentiated 3T3-L1 adipocytes fasted for 12 h and treated with the indicated concentrations of insulin or RF-409 for 10 min. **(B)** Quantification of immunoblot data shown in (A), fit by nonlinear regression. Phosphorylation levels were normalized to total protein and expressed relative to the response to 100 nM insulin. Data are presented as mean ± SEM; *n* = 3 independent experiments. **(C)** Representative immunoblot of C2C12-IR cells fasted for 4 h and treated with insulin or RF-409 (10 nM) for 5 min, followed by ligand washout and incubation for the indicated time points.

**Fig. S2.**
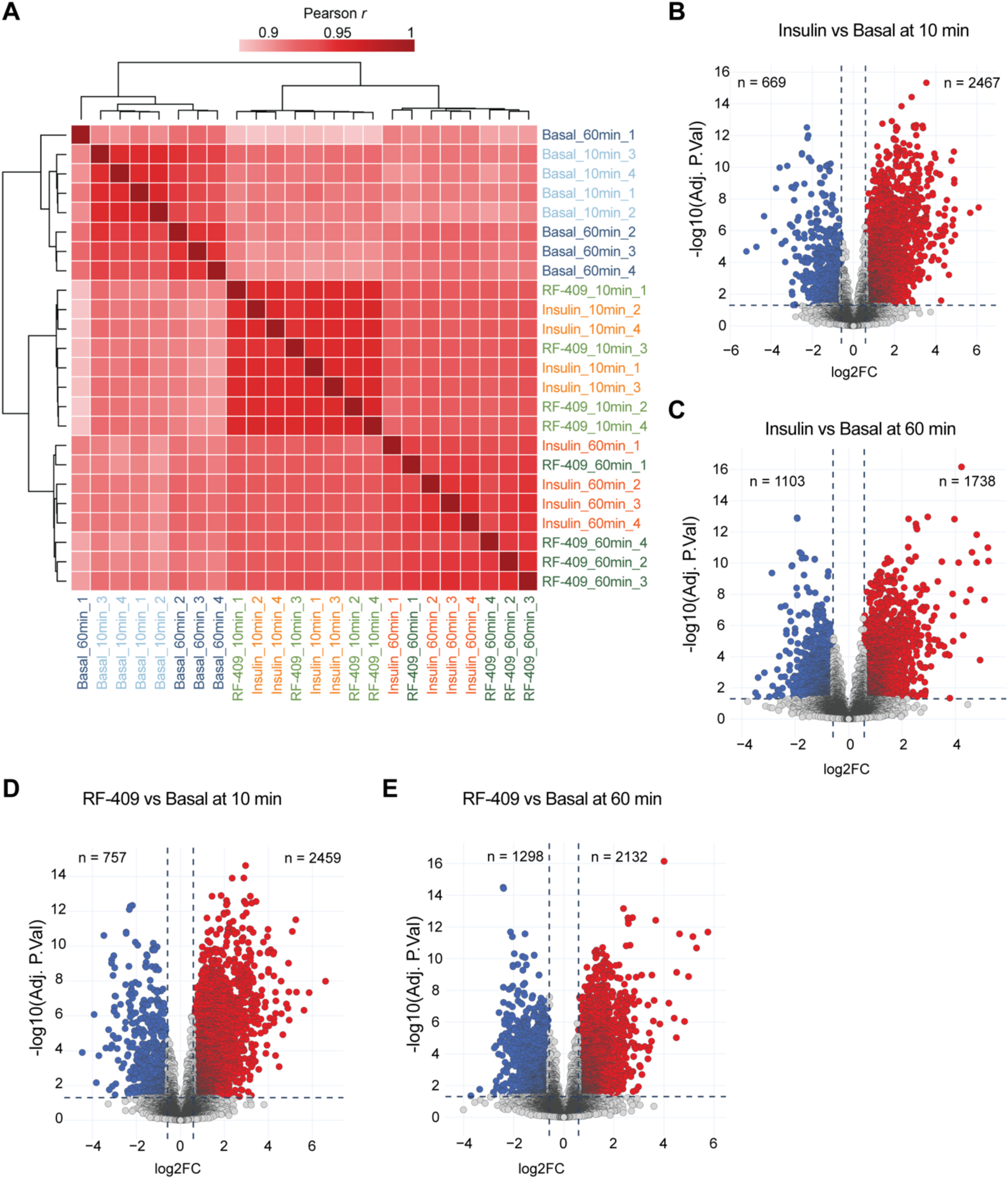
Phosphoproteomic signatures of insulin and RF-409. **(A)** Hierarchically clustered heatmap of phosphoproteomic profiles across basal, insulin-, and RF-409–treated conditions at 10 and 60 min. **(B–E)** Volcano plots showing phosphosites significantly regulated after 10 or 60 min of insulin or RF-409 treatment relative to basal conditions.

**Fig. S3.**
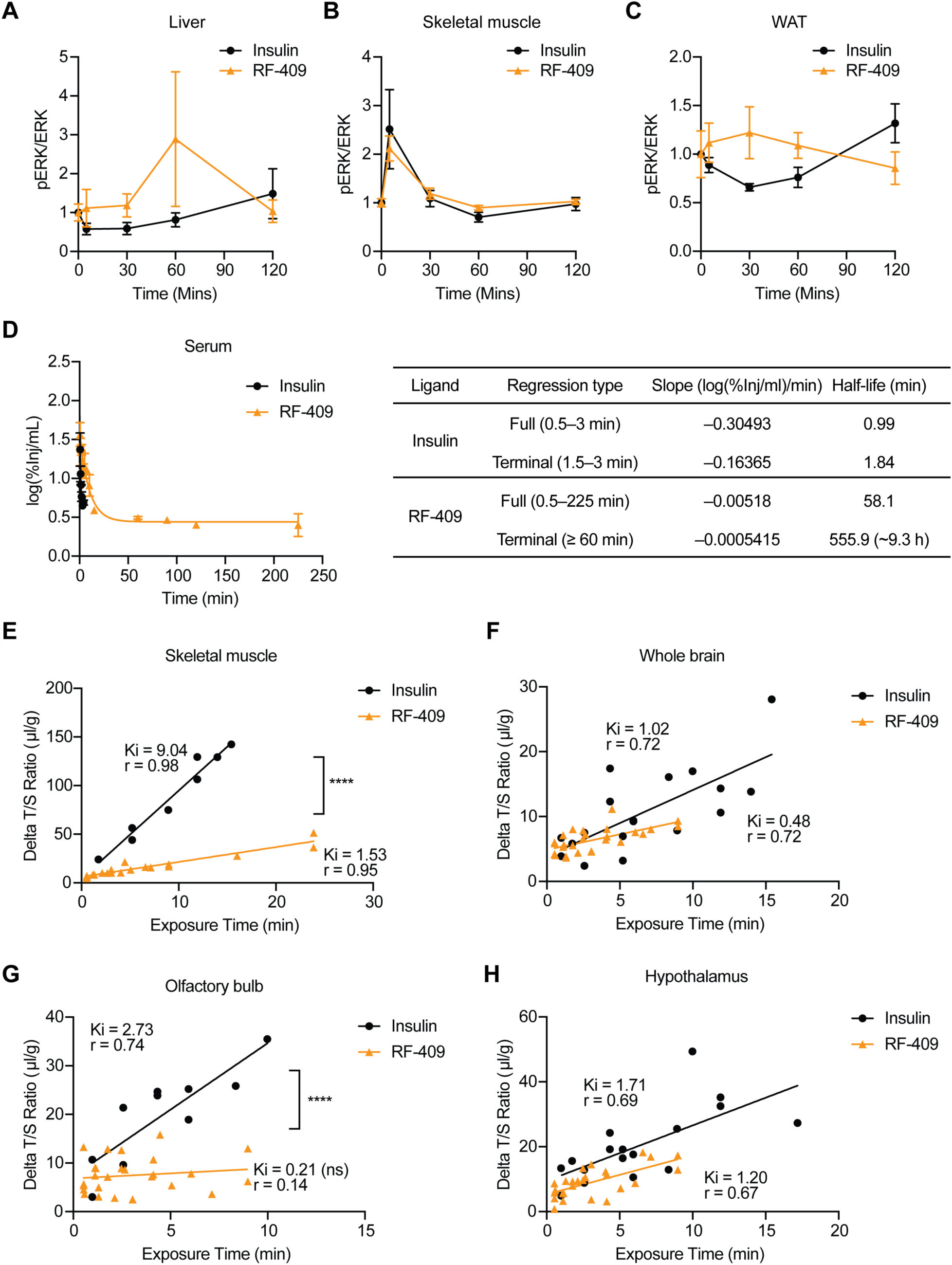
RF-409 exhibits extended stability in vivo and limited tissue penetration. **(A–C)** Pharmacodynamic analysis of ERK phosphorylation (pERK) in liver (A), skeletal muscle (B), and epididymal white adipose tissue (eWAT) (C) following administration of insulin or RF-409, as described in Fig. 3A. **(D)** Extended pharmacokinetic analysis of insulin and RF-409 in serum, with summary parameters. **(E–H)** Tissue/serum ratios of insulin and RF-409 in skeletal muscle (E), whole brain (F), olfactory bulb (G), and hypothalamus (H) are plotted against exposure time. The slopes of the lines for each linear regression represent the unidirectional influx rate (Ki) in units of µl/g-min while the correlation coefficient is represented by ‘r’.

**Fig. S4.**
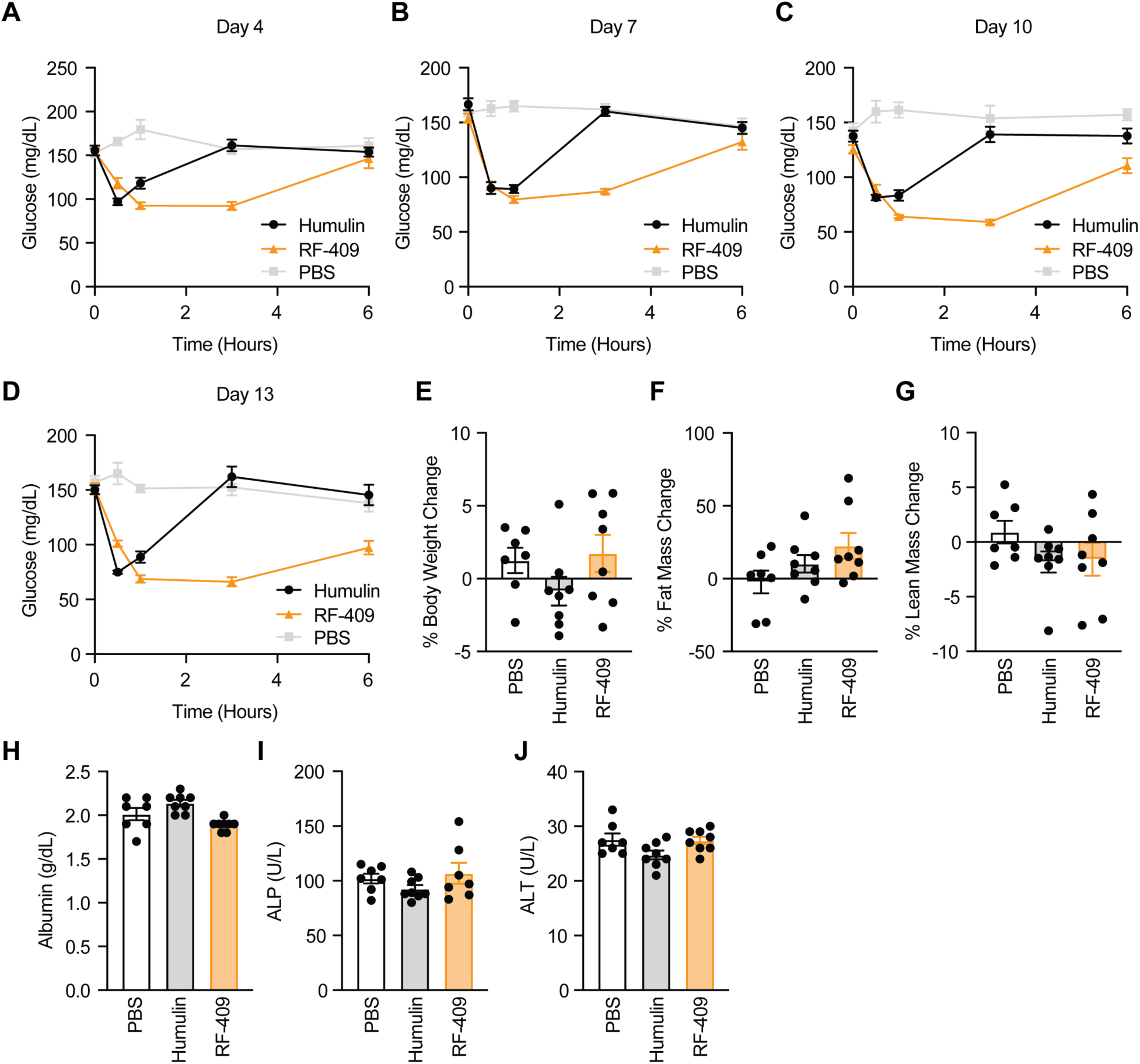
RF-409 produces sustained glucose-lowering effects in vivo. **(A–D)** Ad libitum tolerance tests performed on treatment days 4 (A), 7 (B), 10 (C), and 13 (D) in mice treated with insulin or RF-409, as described in Fig. 3G. **(E)** Percent change in body weight over the treatment period. **(F)** Percent change in fat mass. **(G)** Percent change in lean mass. **(H)** Plasma albumin levels. **(I)** Plasma alkaline phosphatase (ALP) levels. **(J)** Plasma alanine aminotransferase (ALT) levels. Data are presented as mean ± SEM; *n* = 8 (RF-409 and Humulin) and n = 7 (PBS).

**Fig. S5.**
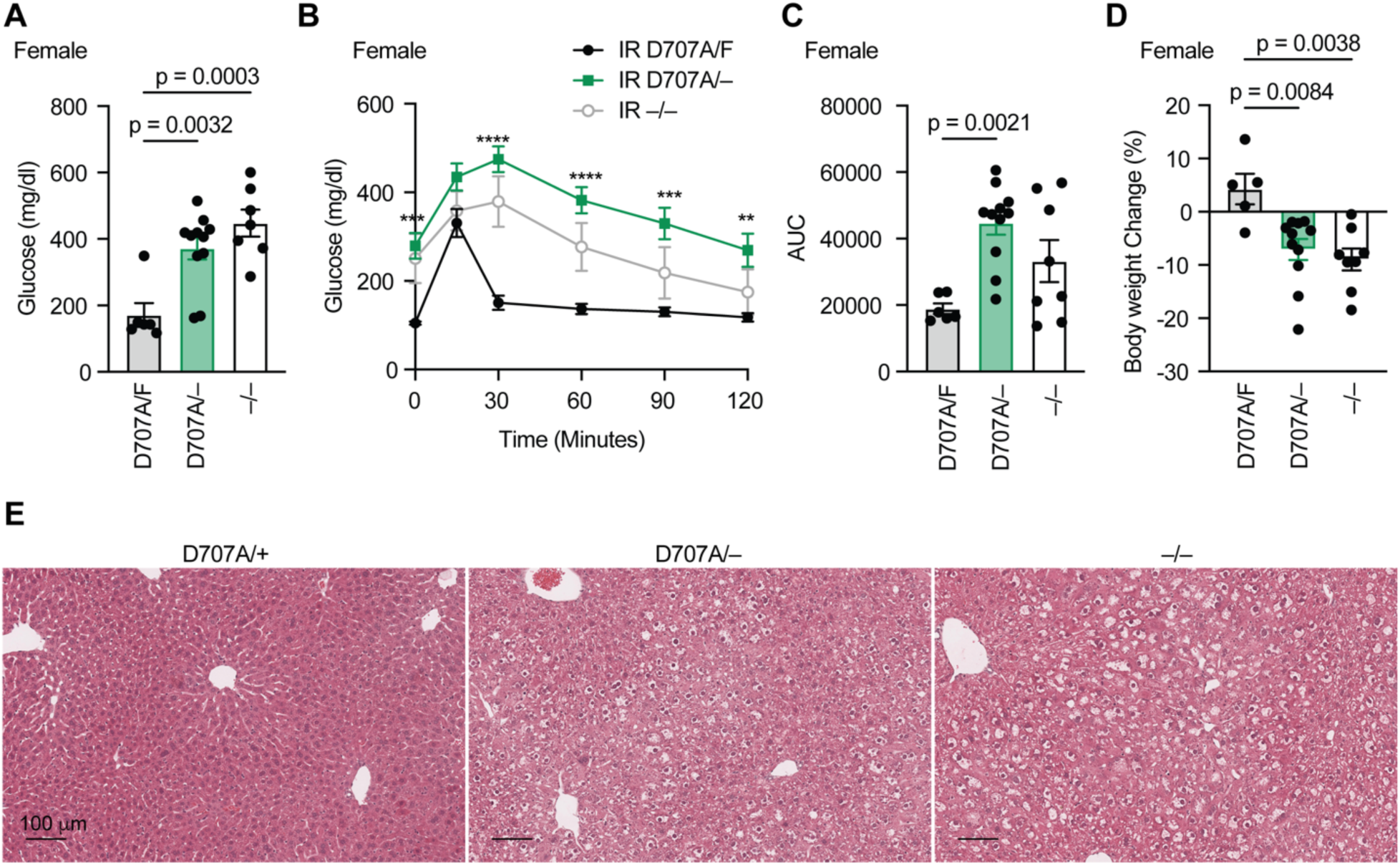
Female and male IR D707A mice exhibit severe insulin resistance. **(A)** Ad libitum blood glucose levels. D707A/F, *n* = 6; D707A/–, *n* = 11; –/–, *n* = 7. Data are presented as mean ± SEM; one-way ANOVA. **(B)** Glucose tolerance test (GTT). Data are presented as mean ± SEM; two-way ANOVA; *n* = 8 mice per group; ****p < 0.0001. **(C)** Area under the curve (AUC) analysis of the GTT shown in (B). Data are presented as mean ± SEM; one-way ANOVA. **(D)** Percent change in body weight between pre-tamoxifen injection and the final day of the experiment. D707A/F, *n* = 5; D707A/–, *n* = 11; –/–, *n* = 8. Data are presented as mean ± SEM; one-way ANOVA. **(E)** Representative H&E staining of male liver tissue.

**Fig. S6.**
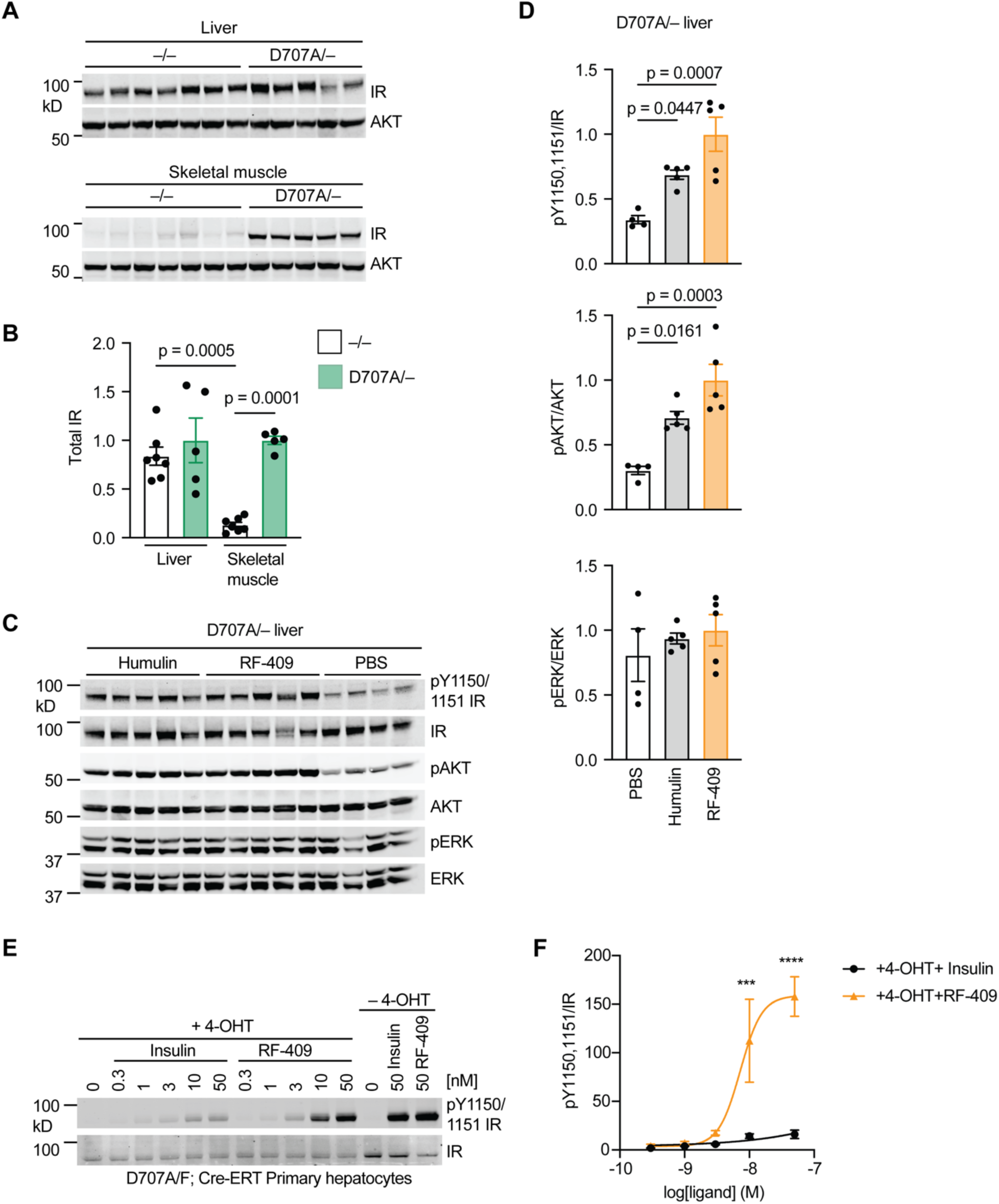
RF-409 activates IR D707A in primary hepatocytes. **(A)** Representative immunoblot of IR protein levels in liver and skeletal muscle from IR D707A/– and IR –/– mice following a 4 h fast. **(B)** Quantification of immunoblot data shown in (A). Data are presented as mean ± SEM; IR D707A/-, *n* = 5; IR –/–, *n* = 7; one-way ANOVA. **(C)** Representative immunoblot of insulin receptor (IR) autophosphorylation, AKT phosphorylation (pAKT), and ERK phosphorylation (pERK) in liver from IR D707A/– mice. Mice were fasted for 4 h and injected with PBS, Humulin (6 nmol/mouse), or RF-409 via the inferior vena cava (IVC). Liver tissue was collected 3 min after injection. **(D)** Quantification of immunoblot data shown in (C). Data are presented as mean ± SEM; PBS, *n* = 4; Humulin, *n* = 5; RF-409, *n* = 5; one-way ANOVA. **(E)** Representative immunoblot of IR autophosphorylation in primary hepatocytes isolated from IR D707A/F;Cre-ERT mice. Cells were treated with 4-hydroxytamoxifen (4-OHT) to delete the floxed WT IR allele and generate IR D707A/– hepatocytes, followed by stimulation with insulin or RF-409 at the indicated concentrations for 10 min. **(F)** Quantification of immunoblot data shown in (E). Phosphorylation levels were normalized to total protein and expressed relative to the response to 100 nM insulin. Data are presented as mean ± SEM; *n* = 3 independent experiments; ****p < 0.0001.

**Fig. S7.**
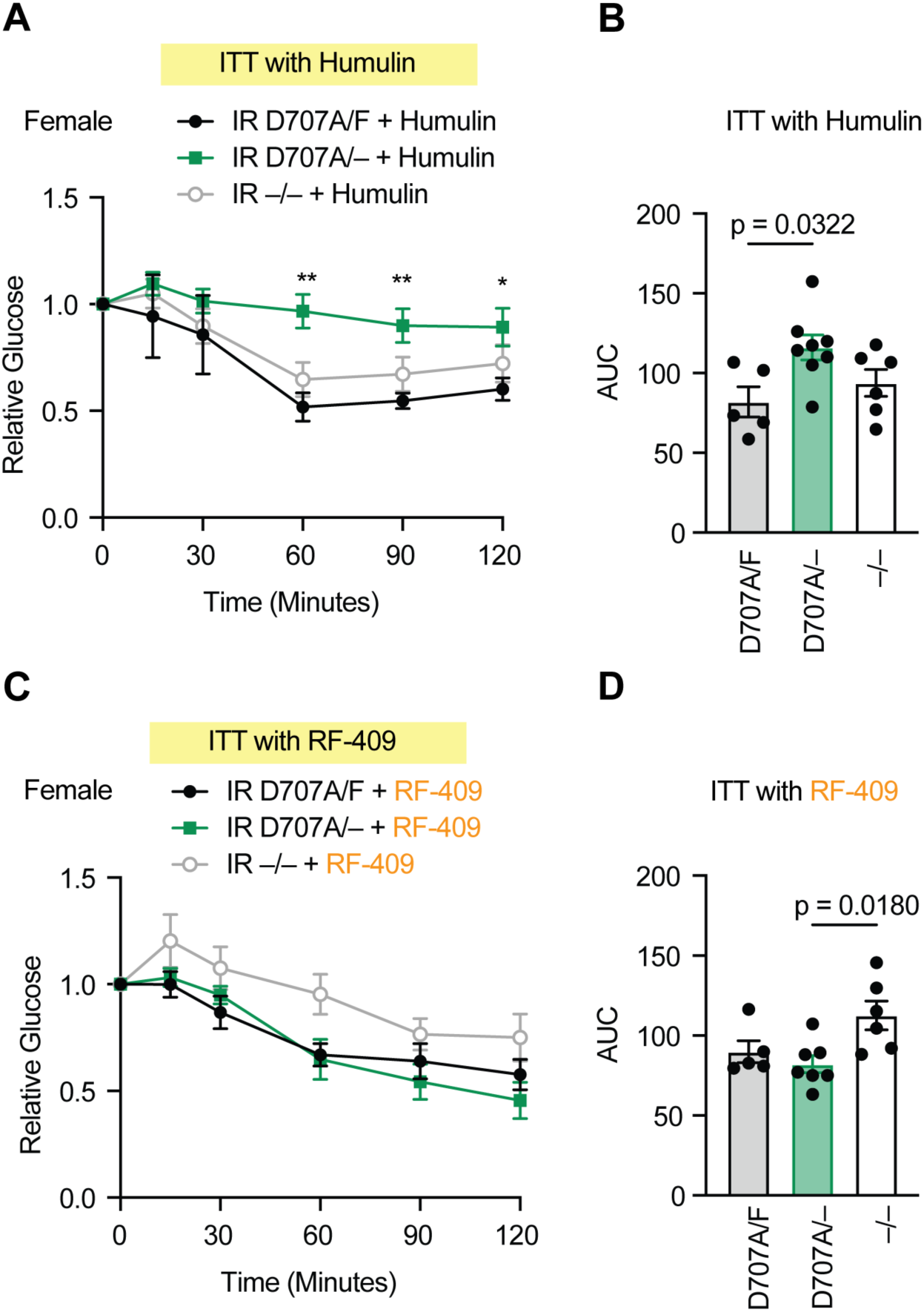
RF-409, but not insulin, lowers blood glucose levels in female IR D707A mice. **(A)** Insulin tolerance test (ITT) following Humulin administration in female mice. IR D707A/F, *n* = 8; IR D707A/–, *n* = 12; IR –/–, *n* = 9. Data are presented as mean ± SEM; two-way ANOVA; ***p < 0.001, ****p < 0.0001 (IR D707A/F vs. IR D707A/–). **(B)** Area under the curve (AUC) analysis of the ITT shown in (A). Data are presented as mean ± SEM; one-way ANOVA. **(C)** Insulin tolerance test (ITT) following RF-409 administration in female mice. IR D707A/F, *n* = 9; IR D707A/–, *n* = 11; IR –/–, *n* = 9. Data are presented as mean ± SEM; two-way ANOVA; *p < 0.05, **p < 0.01, ***p < 0.001 (IR D707A/F vs. IR D707A/–). **(D)** Area under the curve (AUC) analysis of the ITT shown in (C). Data are presented as mean ± SEM; one-way ANOVA.

